# Flap-enabled next-generation capture (FENGC): precision targeted single-molecule profiling of epigenetic heterogeneity, chromatin dynamics, and genetic variation

**DOI:** 10.1101/2022.11.08.515732

**Authors:** Mingqi Zhou, Nancy H. Nabilsi, Anqi Wang, Marie-Pierre L. Gauthier, Kevin O. Murray, Hassan Azari, William S. Owens, Jeremy R. B. Newman, Francisco J. Pardo-Palacios, Ana Conesa, Alberto Riva, Thomas L. Clanton, Brent A. Reynolds, Patrick Concannon, Jason O. Brant, Rhonda Bacher, Michael P. Kladde

## Abstract

Targeted sequencing is an increasingly sought technology. Available methods, however, are often costly and yield high proportions of off-target reads. Here, we present FENGC, a scalable, multiplexed method in which target sequences are assembled into 5′ flaps for precise excision by flap endonuclease. Recovery of length-matched sequences, amplification with universal primers, and exonucleolytic removal of non-targeted genomic regions mitigate amplification biases and consistently yield ≥ 80% on-target sequencing. Furthermore, optimized sequential reagent addition and purifications minimize sample loss and facilitate rapid processing of sub-microgram quantities of DNA for detection of genetic variants and DNA methylation. Treatment of cultured human glioblastoma cells and primary murine monocytes with GC methyltransferase followed by FENGC and high-coverage enzymatic methyl sequencing provides single-molecule, long-read detection of differential endogenous CG methylation, dynamic nucleosome repositioning, and transcription factor binding. FENGC provides a versatile and cost-effective platform for targeted sequence enrichment for analysis of genetic and/or epigenetic heterogeneity.

## Introduction

Multiplexed enrichment of target sequences for next-generation sequencing (NGS) has diverse applications in basic science and medicine^1^. The reduced sample complexity imparts substantial NGS cost savings, as well as higher per base coverages that improve data quality and detection of rare variants. Available methods, however, generally interrogate either genetic or epigenetic signatures, require expensive reagents, have limited flexibility regarding target design, and exhibit high off-target rates. Furthermore, sequence recovery is usually poor due, necessitating high amounts of input DNA. In addition, most enrichment techniques capture relatively short fragments with wide length distributions from which the shorter fragments are preferentially amplified.

Hybridization-based target enrichment using tiled DNA or RNA oligonucleotide ‘baits’ is widely used for targeted applications such as DNA methylation analysis^2^. Although tiling improves coverage uniformity, pools of overlapping, long (50-170 nt), and biotinylated oligonucleotides^3^ are cost prohibitive to many laboratories. Hybridization enrichment also suffers from long incubation times, high off-target rates, and inefficient recovery of target sequences^4^.

Random fragmentation of input DNA further contributes to off-target sequencing, as well as amplification and sequencing biases. Lastly, hybridization-based capture that yields short reads precludes phasing of epigenetic and genetic information.

Clustered, regularly interspaced short palindromic repeats (CRISPR)-associated (Cas) enzymes, which utilize short RNAs to direct site-specific DNA cleavage^5^, are increasingly enlisted for sequence enrichment *in vitro*. However, for multiplexed enrichment of shorter sequences with CRISPR-Cas9 (refs. 6-9), the requirement for a 2-bp protospacer adjacent motif constrains target design and thus length matching, resulting in amplification and sequencing biases. This drawback is in part circumvented by direct sequencing of captured megabase-length fragments with Nanopore technology^10^; however, for clinical applications, the requirement for large amounts of DNA is problematic. In addition, accurate single-molecule detection of 5-methylcytosine (5mC) as opposed to deriving a population averaged view remains technically challenging. Lastly, off-target cutting^11,12^ and the expenses of recombinant Cas enzymes and guide RNAs constitute further barriers to scaling for many laboratories.

Bisulfite patch PCR^13^, optionally combined with the methyltransferase accessibility protocol for individual templates (MAPit), termed MAPit-patch^14^, are additional methods for targeted genetic and epigenetic surveys. These methods, however, rely on restriction endonucleases, which severely limit target selection design and generate fragments with broad length distributions that introduce substantial amplification and sequencing biases.

Here, we have developed and applied flap-enabled next-generation capture (FENGC), a versatile, rapid, cost-effective, and scalable method for multiplexed sequence enrichment. FENGC enlists flap endonuclease (FEN) to excise target sequences reconstituted into 5′ DNA flaps to achieve high ratios of on-to off-target enrichment. Programmable, nucleotide-specific cleavage impart unparalleled versatility in target design and capture, enabling matching sequence lengths to mitigate PCR bias and further improve on-target sequencing frequencies. FENGC of genomic DNA (gDNA) from cultured and primary cells treated with GpC methyltransferase followed by enzymatic methyl sequencing (EM-seq) and high-fidelity, long-read sequencing reproducibly localizes nucleosomes, transcription factor (TF) binding, and endogenous CG methylation on contiguous single molecules. Furthermore, the achieved high sequencing coverages facilitate detection of low-level, but significant differences, in chromatin accessibility and DNA methylation, as well as dynamic nucleosome repositioning as a driver of nucleosome-free region (NFR) length. FENGC without deamination provides high-confidence identification of single-nucleotide polymorphisms (SNPs) and insertions/deletions (indels).

## Results

### FENGC, a novel, cost-effective method for on-target sequence enrichment in genetic and epigenetic studies

To establish FENGC, we retained advantages of multiplexed DNA strand capture and amplification afforded by patch ligation PCR^13,14^, but eliminated the requirement for restriction enzymes. To do so, we leveraged the activities of thermostable *Thermococcus* 9° N flap endonuclease 1 (FEN1) and the FEN domain of Taq DNA polymerase I, which tightly bind and efficiently cut double flaps (unpaired 5’ flap and 1-nt 3’ flap) with exquisite nucleotide-level precision and uniformity^15-19^.

In a multiplexed reaction, FENGC enlists a pair of target-specific flap adapters to assemble both ends of each target sequence into a double flap (Fig. 1a; Supplementary Fig. 1). Following specific cleavage by FEN1 or Taq, target sequences are patch ligated to universal primers, one of which protects against subsequent exonuclease digestion. The enriched sequences are purified and amplified by standard PCR for genotyping or, alternatively, subjected to bisulfite- or enzyme-mediated deamination of C to U prior to amplification (methyl-PCR) for base-level detection of 5mC. Shorter library products can be barcoded and sequenced as described previously^14^. Herein, longer PCR and methyl-PCR products were examined by long-read, high-fidelity (Hi-Fi; >Q30) circular consensus sequencing (CCS; Pacific Biosciences, PacBio).

**Fig. 1.**
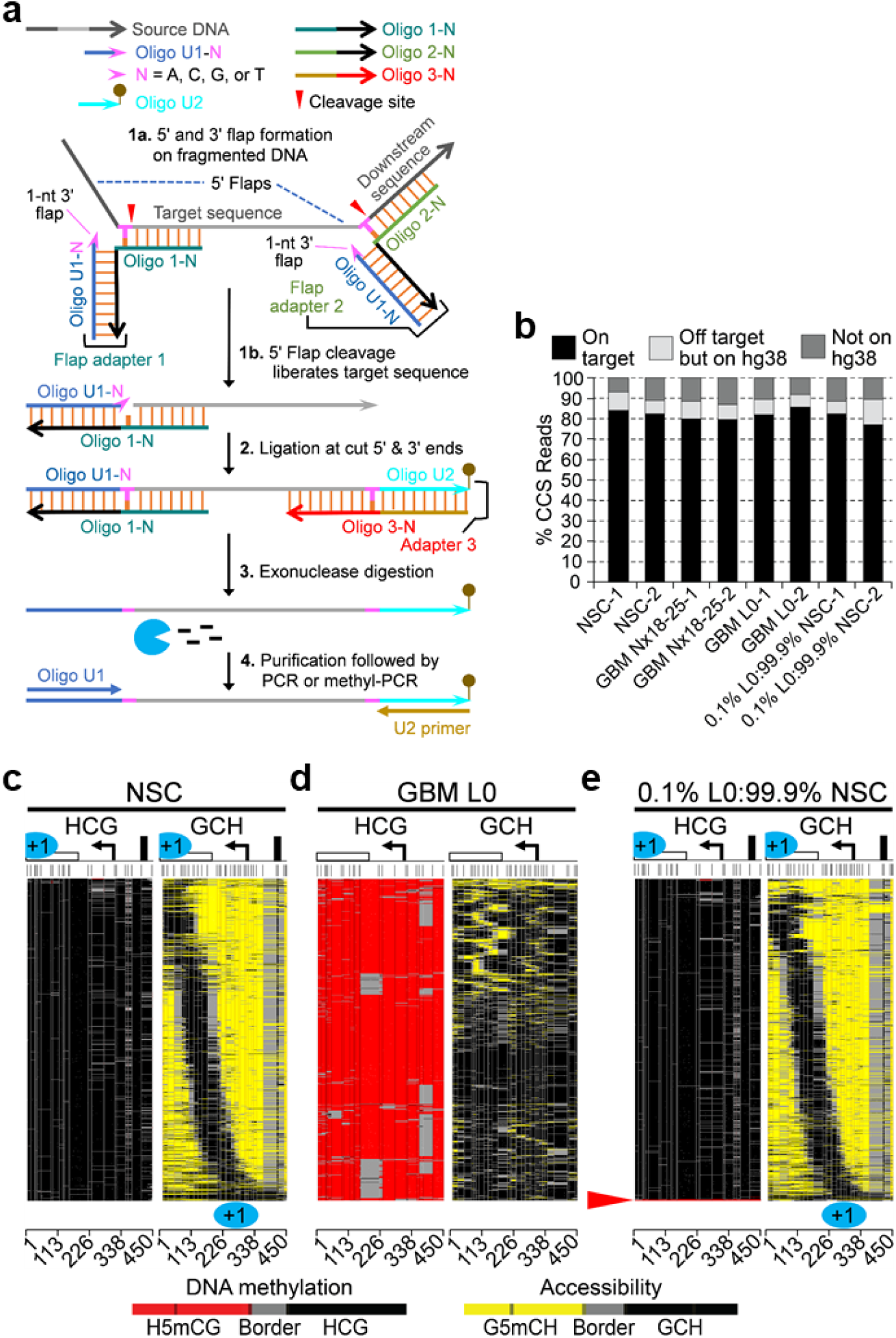
FEN activity-dependent target sequence enrichment yields high percentages of on-target reads, profiles epigenetic heterogeneity, and detects low-frequency epialleles. **a**, FENGC procedure. Mildly fragmented gDNA is hybridized to a pair of locus-specific flap adapters, reconstituting a double flap (5’ and 1-nt 3’ flap) at both ends of each target sequence (step 1a). To form 1-nt 3’ flaps, nucleotides (emphasized in pink) 5’ of the scissile phosphodiesters (red arrowheads) ‘overlap’ and thus displace the 3’-terminal nucleotide of oligo U1-A, C, G, or T (oligo 1-N). Specific cleavage by FEN activity of FEN1 or Taq DNA polymerase I releases single-stranded target strands with ligatable ends, i.e., 5’ phosphate downstream of a 1-nt gap and a 3’-hydroxyl (step 1b). The gap is filled by the 1-nt 3’ flap, which ligates to the cut 5’ end (step 2). A nested 3’-end-specific adapter 3 is added to position oligo 2 for ligation to the cut 3’ end. Covalent modification of oligo U2 (lollipop) protects against 3’ to 5’ exonuclease degradation, dramatically enriching target sequences (step 3). All reagent addition to this point is sequential in a single tube, avoiding loss of target sequences. Enriched targets are purified and amplified with either standard PCR or methyl-PCR to construct NGS libraries (step 4). **b**, Proportions of Hi-Fi EM-seq CCS reads for MAPit-FENGC of 119 promoter targets mapping to or off the human genome. Suffixes denote the two independent biological replicates for the indicated sequenced libraries. **c-e**, Methylscaper heatmaps of HCG and GCH methylation of single molecules generated from filtered EM-seq CCS reads aligning to *EPM2AIP1* (+307 to **–** 143) in the combined replicates of NSC (3,460 reads) (**c**), GBM L0 (3,889 reads) (**d**), and 0.1% L0 gDNA spiked into NSC gDNA (1,781 reads) (**e**). In all such pairs of plots, the patterns of HCG methylation (left; consecutive H5mCG colored red) and GCH accessibility (right; consecutive G5mCH colored yellow) on each molecule or epiallele shown in the same top-to-bottom order. Two or more consecutively unmethylated sites are connected by black, gray connects the borders between methylated and unmethylated sites, and white indicates missing or unaligned sequences. Arrow, TSS; white rectangle, part of *EPM2AIP1* protein coding exon; blue partial and full ovals, footprinted +1 nucleosomes; black rectangle, footprinted TF. Vertical ticks and lines across panels mark respective HCG and GCH motifs. In these and subsequent plots, all features and intervening distances are drawn to scale. A proportions test in R indicated that the proportion of 10 CG-hypermethylated of 1,781 total epialleles detected from the duplicate samples in **e** is at least or greater than 0.1% (*P* = 3.59E**–** 09). Base pair position along each target is indicated at bottom.

### FENGC optimization and validation

We first determined that *Thermococcus* 9° N FEN1 cut an oligo (proxy target) assembled into a 5’ flap more efficiently than Taq (Supplementary Fig. 2; Supplementary Table 1, Sheet 1). Next, we tested the effect of DNA methylation and changing the base of the ‘overlapped’ nucleotide on FEN1 activity (Supplementary Fig. 3). In this experiment, a pair of unmethylated and methylated 80-mers was reconstituted into four double flaps with A, C, G, or T both 5’ of the scissile phosphodiester and as a 1-nt 3’ flap (Supplementary Fig. 3a; Supplementary Table 1, Sheet 1). Five C or 5mC residues in the two 80-mers reside near the 5’ flap cleavage sites, locations contacted by human FEN1 co-crystallized with a double flap (5’ flap and 1-nt 3’ flap)^19^. Importantly, the use of unmethylated flap adapters parallels FENGC enrichment of methylated targets. Equally efficient cleavage of all four double flaps ± nearby 5mC (Supplementary Fig. 3b,c) suggests FENGC is compatible with DNA methylation analyses, including MAPit methylation footprinting^20^, also known as NOMe-seq^21^.

We first examined the ability of FENGC to enrich single-copy genomic sequences from gDNA of HCT116 colon cancer cells, as well as the method’s suitability for bisulfite sequencing. Flap adapters were designed to excise 11 promoters, including 9 from DNA repair genes, *EMP2AIP1* (transcribed divergently from *MLH1*), and *HSPA5* as an open chromatin positive control (Supplementary Table 1, Sheets 2 and 3). Sequences had mean lengths of 298 nt (range 282-315 nt) and 445 nt (range 430-452 nt), with both sets anchored at the 5’ end (Supplementary Table 2, Sheets 1 and 2). The ∼ 300-nt and ∼ 450-nt targets were most likely cut and ligated to oligos U1-T and U2 (totaling 47 nt), given amplification of the expected ∼ 350 bp and ∼ 500 bp products, respectively, dependent on the FEN activity of Taq and FEN1 (Supplementary Fig. 4a, compare lane 1 with 2 and 3 with 4; Supplementary Fig. 4b, compare lane 1 with 2 and 4 with 5). Superior amplification yield was reproducibly observed with FEN1 vs. Taq (Supplementary Fig. 4a, compare lane 2 with 4). As expected, FENGC product yields were reduced with longer amplicons and bisulfite (BS) treatment, which hydrolyzes DNA (Supplementary Fig. 4b, compare lane 2 with 5 and lane 3 with 6, respectively). Therefore, detection of the ∼500 bp BS-PCR product required 4 μg gDNA input for this FENGC reaction that targeted only 11 loci (compare lane 6 with 7).

### MAPit-FENGC, a versatile assay for targeted genetic and epigenetic analyses

Given the successful post-capture bisulfite conversion and amplification of target sequences, we combined MAPit methylation footprinting and FENGC. The resulting MAPit-FENGC method was tested on biological duplicates of human cell line L0 derived from a glioblastoma (GBM) tumor of the mesenchymal subtype^22^. Accessible GC sites in chromatin of permeabilized cells were marked by incubation with GpC methyltransferase M.CviPI and methyl donor *S*-adenosyl-*L*-methionine (Methods). FENGC with Taq or FEN1 was used to target the same 11 promoter sequences of ∼ 300 nt and ∼ 450 nt, followed by standard PCR or BS-PCR and sequencing using PacBio Hi-Fi long-read CCS (Methods).

In this MAPit-FENGC pilot, ≥ 1 CCS read aligned to at least 9 of the 11 target promoters (82%) in all 12 reactions (Supplementary Table 3). Zero to few reads were detected for *MSH3* and *MSH6*, possibly due to their high GC content, typical of NGS and patch PCR methods^13,14^. For the remaining 9 targets, FENGC with both Taq and FEN1 using standard PCR achieved

≥ 25x coverage in the reads pooled from both samples for each condition. For FENGC with FEN1 and BS-PCR, ≥ 150x coverage was observed in the pooled reads for the same 6 targets in both the ∼ 300-nt and ∼ 450-nt sequences. For these 6 targets, methylation levels at HCG (H5mCG) and GCH (G5mCH), i.e., excluding GCG (overlap between GC and CG), were highly correlated between both biological replicates and between the shared 5’ ends of the enriched ∼ 300-nt and ∼ 450-nt sequences (Supplementary Fig. 5; *r* ≥ 0.92).

To increase the sensitivity of FENGC targeted DNA methylation analysis, bisulfite was substituted with EM-seq, a nondestructive, enzymatic means to convert C to U ^23,24^. For this analysis, the number of target promoters was scaled from 11 to 119, adding the promoters of 74 genes with Gene Ontology term “DNA repair,” 41 cancer-associated genes, and 3 promoters with open chromatin (Supplementary Table 1, Sheet 3; Supplementary Table 2, Sheet 2). For *FAT4*, oligo 2-T was designed to not cut the target 3’ end, therefore providing a negative control. This 119-target panel (range 430-452 nt) was used for MAPit-FENGC of different amounts of gDNA from M.CviPI-treated L0 cells. Subsequent EM-seq and amplification (EM-PCR) produced the specific, expected ∼ 500 bp product, including 47 bp universal primers, from as little as 50 ng gDNA (Supplementary Fig. 6). By contrast, bisulfite conversion required 80x more gDNA (4 μg) to detect an amplicon (Supplementary Fig. 4b).

Next, MAPit-FENGC was conducted on two independent cultures of GBM line L0, GBM Nx18-25 (classical subtype), and for comparison, non-cancerous human fetal neural stem cells (NSC)^22^. GDNA from these M.CviPI-treated lines and two 0.1% L0 gDNA:99.9% NSC gDNA mixtures was processed in parallel by FENGC with the 119-promoter panel followed by EM-seq conversion. The ranges of PacBio Hi-Fi CCS reads that aligned to the target references of the hg38 reference genome, off-target but on hg38, and off hg38 from the 8 libraries were 77-86%, 6.6-9.9% (excluding an outlying L0:NSC mixture at 14%), and 6.9-13% (Fig. 1b; Supplementary Table 4, Sheet 1). These on-target percentages out-perform single-molecule footprinting using tiled RNA baits to capture smaller fragments of 200-300 bp^25^. As expected, zero reads from all eight libraries aligned to negative control locus *FAT4* (Supplementary Table 5). MAPit-FENGC detected 105-111 targets (89-94%, excluding *FAT4*) with ≥ 1 read in the combined biological duplicates for each sample (Supplementary Table 4, Sheet 1). To avoid multiple alignments to gene homologs, reads were further filtered for ≥ 95% coverage of each reference sequence length. These filtered reads aligned uniquely to 92-105 targets (78-89%; Supplementary Table 4, Sheet 1; Supplementary Table 5), and read number *versus* GC content showed a modest negative fit (Supplementary Fig. 7).

For the fractions of H5mCG (DNA methylation) and G5mCH (chromatin accessibility), all correlation coefficients between the independent biological duplicates of each sequenced library were > 0.91 (Supplementary Fig. 8). In addition, the proportions of H5mCG and G5mCH were highly correlated between the EM- and BS-PCR data sets, indicating that EM-seq can be substituted for BS-seq in MAPit-FENGC (Supplementary Fig. 9; *r* > 0.93).

### MAPit-FENGC with EM-seq informs epigenetic landscapes and efficiently detects differential DNA methylation and accessibility in minor subpopulations of cells

Among the 119 loci profiled by MAPit-FENGC, the *EPM2AIP1* promoter exemplifies marked epigenetic reprogramming in GBM (Fig. 1c-d). To reveal methylation patterns, molecules were hierarchically clustered based on the first principal component of H5mCG and G5mCH levels by methylscaper^26^. In non-cancerous NSC, the promoter was predominantly unmethylated and highly accessible to M.CviPI, except for a +1 nucleosome (partial and full blue ovals; expected ∼ 150-bp protections), which occupied a continuum of positions in different cells across the sample population, thereby lengthening or shortening NFRs (Fig. 1c, right panel). These accessible, variable-length NFRs were punctuated by a short footprint of fixed location (black rectangle), indicative of strong sequence-specific TF binding. By contrast to NSC, in GBM L0, the *EPM2AIP1* promoter was hypermethylated at HCGs (Fig. 1c,d, compare left panels) and closed, except for relatively short, accessible linkers within randomly positioned nucleosomes (Fig. 1c,d, compare right panels).

Challenging the sensitivity of MAPit-FENGC, analysis of the duplicate mixtures of 0.1% L0 gDNA:99.9%NSC gDNA revealed detection of a significant proportion of hypermethylated *EPM2AIP1* molecules, likely originating from the 0.1% L0 spike-in (Fig. 1e, red arrowhead; *P* = 3.59E**–** 09). The chromatin states of the other molecules resembled those of NSC (Fig. 1d, right). Together, these data show that FENGC maps epigenetic states reproducibly, quantitatively, and with high sensitivity, detecting as few as 0.1% hypermethylated epialleles.

The ability of MAPit-FENGC to detect differential epigenetic landscapes was tested further by assaying the same 119-target panel from two independent cultures of non-cancerous NSC and GBM Nx18-25. Epigenetic differences between NSC and GBM were assessed for 54 targets with ≥ 50 total CCS reads in both replicates, but no less than 22 reads in either replicate (Supplementary Table 6). Using a significance criterion of *P* < .05 and differential levels of either H5mCG or G5mCH of ≥ 0.05, ≥0.10, ≥ 0.20, and ≥0.45, 57% (31/54), 24% (13/54), 13% (7/54), and 7.4% (4/54) of promoters exhibited candidate epigenetic differences, respectively (Supplementary Table 6). The quantitative nature of these results is supported by the highly correlated (*r* > 0.91) H5mCG and G5mCH levels between replicates (Supplementary Fig. 8). In addition, 733 of 735 total *MSH5* promoter copies (all 4 samples) had ≥10 G5mCH residues (Extended Data Fig. 1). Such high accessibility rules out incomplete cell permeabilization and limiting M.CviPI activity as possible bases for differential accessibilities, either between different loci or molecules from a given locus in the same or different samples.

To illustrate quantitative detection of epigenetic differences between NSC and GBM Nx18-25, three representative promoters were chosen and correlated with transcript levels (Fig. 2; Extended Data Fig. 2). *CD44* encodes a transmembrane glycoprotein with functions in cell adhesion, proliferation, and apoptosis^27,28^. *CD44* expression is also a marker of astrocyte-restricted precursors^29^. In particular, *CD44* overexpression in GBM tissue has been linked to the mesenchymal subtype^30,31^ and GBM cancer stem cells^32,33^. MAPit-FENGC of NSC showed methylation of two to three HCGs upstream of the *CD44* TSS (Fig. 2a, panel 1). Notably, the longest NFR on most of the NSC molecules occupied diverse positions, which coincided with movement of the −1 nucleosome across the entire assayed region (Fig. 2a, panel 2). In striking contrast to NSC, in GBM Nx18-25, reproducible, pronounced, and variable-length expansion of H5mCG was observed (Fig. 2a, panel 1 *vs*. 3; Fig. 2b, left, *P* < .0001). The increased H5mCG in Nx18-25 (compared with NSC) correlated with marked, significant reductions in accessibility (Fig. 2a, panel 2 *vs*. 4; Fig. 2b, right, *P* < .0001), cumulative NFR length (Fig. 2c, *P* = 2.2E-16), and transcript abundance (Fig. 2d, *P* < .01).

**Fig. 2.**
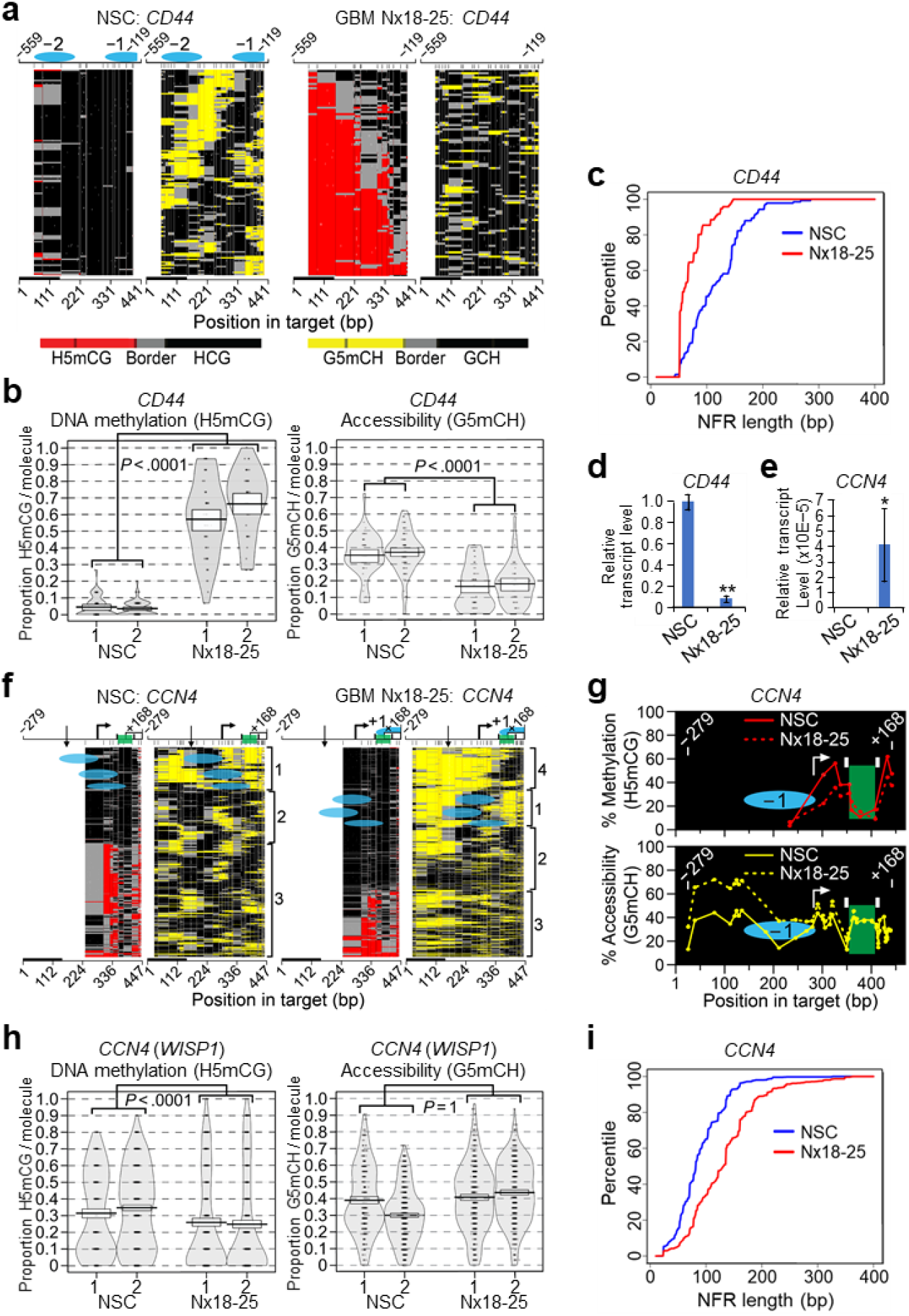
Targeted MAPit-FENGC detects differential epigenetic configurations that correlate with gene expression. **a-c**, Filtered EM-seq CCS reads aligning to *CD44* from human NSC (*n* = 2; 141 combined reads) and GBM Nx18-25 (*n* = 2; 96 combined reads) plotted by methylscaper as clustered single-molecule methylation heatmaps (**a**), violin plots of the per molecule proportion of H5mCG (endogenous DNA methylation, left) and G5mCH (accessibility, right) (**b**), and cumulative distribution function of NFR length (mean = 118 bp in NSC and 72.0 bp in Nx18-25, *P* = 2.2E**–** 16) (**c**). **d,e**, Relative expression of *CD44* (*n* = 3, mean ± SD; **, *P* < .01) (**d**) and *CCN4* (*n* = 3, mean ± SD; paired Student’s *t* test, two sided, *, *P* < .05) (**e**). **f-i**, Filtered EM-seq CCS reads aligning to *CCN4* from human NSC (*n* = 2; 1,116 combined reads) and GBM Nx18-25 (*n* = 2; 1,143 combined reads; Supplementary Table 5) plotted by methylscaper (**f**), percentages of averaged H5mCG and G5mCH (**g**), violin plots (**h**), and cumulative distribution function of NFR length (mean = 89.4 bp in NSC and 132 bp in Nx18-25, *P* = 2.2E**–** 16) (**h**). Although the difference in H5mCG per molecule between NSC and GBM Nx18-25 in **h** was significant, the difference in accessibility was not, probably due to uncharacteristically high accessibility of molecules bearing H5mCG. Green rectangles in **f,g** demarcate local depletion of H5mCG between two footprinted factors (black rectangles) in cluster 1 molecules. In **f**, a known GA>A indel (dbSNP, rs548251181; straight arrow; plotted in white as it does not align to hg38) present at similar frequencies in both cell types was verified by MAPit-FENGC genotyping with standard PCR amplification (Supplementary Table 7, Indels). White rectangle, coding exon 1 sequence.

CCN4, also known as WNT1-inducible signaling pathway protein 1 (WISP1), contributes to the tumorigenesis and progression of a wide array of human cancers^34-36^. *CCN4* expression is upregulated in GBM compared with normal tissues^37^. Indeed, *CCN4* transcripts were undetectable in NSC and increased dramatically in GBM Nx18-25 (Fig. 2e; *P* < .05). MAPit-FENGC of the *CCN4* promoter from NSC revealed a majority of molecules with variable levels of H5mCG and disorganized nucleosomes (Fig. 2f, panels 1 and 2, respectively; clusters 2 and 3). Molecules in cluster 1, by contrast, exhibited larger spans of accessibility and contained two footprints (black rectangles); however, nucleosomes (blue ovals) occluded the TSS. In terms of peaks and valleys, plots of averaged H5mCG and accessibility were qualitatively similar between NSC and GBM Nx18-25 (Fig. 2g). Quantitatively, however, in GBM, the number of molecules bearing 5mCG declined significantly (Fig. 2f, panel 1 *vs*. 3; Fig. 2g, upper; Fig. 2h, left, *P* < .0001), and NFR length per molecule increased, particularly in newly appearing cluster 4 (Fig. 2f, panel 2 *vs*. 4; Fig. 2i, *P* = 2.2E-16). Overall, the quantitative changes in *CCN4* promoter DNA methylation and accessibility are consistent with the observed marked increase in transcript abundance in GBM (Fig. 2e).

*HIST1H1B* encodes the linker histone protein H1.5, which maintains higher-order chromatin structure, as well as regulates DNA repair and cell proliferation^38,39^. Based on NFR size, position of nucleosome +1, TF binding upstream of the TSS, and DNA hypermethylation, *HIST1H1B* promoter copies from both NSC and GBM Nx18-25 populated seven clusters (Extended Data Fig. 2). Hypermethylated *HIST1H1B* promoter copies increased significantly from 0.6% to 4% in NSC *versus* GBM (Extended Data Fig. 2a, panel 1 *vs*. 3; Extended Data Fig. 2b; Extended Data Fig. 2c, left; Supplementary Table 6, *P* < .0001). In addition, cumulative accessibility, in particular from **–** 80 to +45, decreased significantly in Nx18-25 compared with NSC (Extended Data Fig. 2a, panel 2 *vs*. 4; Extended Data Fig. 2b, bracketed area; Extended Data Fig. 2c, right; Supplementary Table 6, *P* < .0001). However, the modest reduction in accessibility and increase to 4% hypermethylated *HIST1H1B* epialleles in Nx18-25 did not significantly alter bulk transcript abundance (Extended Data Fig. 2d). Taken together, we conclude that MAPit-FENGC efficiently detects prominent as well as low-frequency epigenetic differences, and the profiled epigenetic heterogeneity informs mechanism.

### FENGC detects known and novel genetic alterations

For genotyping, gDNA from the same two independent cultures of M.CviPI-treated NSC and GBM Nx18-25 were processed by FENGC with the 119-promoter panel of ∼ 450 nt (Supplementary Table 2, Sheet 2), without deamination prior to PCR. Hi-Fi CCS reads from the 4 sequenced libraries aligned to the 119 hg38 target references, off-target but on hg38, and off hg38 with ranges of 73-89%, 10-20%, and 0.6-8.9%, respectively (Supplementary Table 4, Sheet 2). Among the 97 targets (82%) with ≥ 1 read in at least one condition, we identified 51 single-nucleotide polymorphisms (SNPs) and 17 indels distributed over 30 promoters (Supplementary Table 7). There were 43 SNPs and 16 indels already recorded in dbSNP (http://www.ncbi.nlm.nih.gov/SNP/). The C-to-A substitution in the *CDH1* promoter at chr16:68737131 (rs16260) was previously identified as a cancer-specific risk factor with clinical significance^40-42^. The SNP A allele was identified within 0% of reads in NSC and 38% of reads in Nx18-25. Eight SNPs and one indel without dbSNP entries were also detected. Reproducible identification of alleles of both known and previously unknown polymorphisms demonstrates the efficacy of FENGC in genotyping.

*Bona fide* epigenetic perturbations, by definition, require no concurrent DNA mutations. Therefore, we further examined if candidate epigenetic differences in targets (Supplementary Table 6) could be attributable at least in part to genetic variation between NSC and GBM Nx18-25 (Supplementary Table 7). A GBM-specific SNP mapping to the *CHEK1* promoter eliminated a CG site, fully accounting for the loss of H5mCG in Nx18-25 *versus* NSC. Allelic differences between NSC and GBM in nine other promoters could in part underpin the identified epigenetic variations. Comparing NSC and GBM, a known indel was represented at similar frequencies in *CCN4* and no allelic variants were detected in *CD44* or *HIST1H1B*, satisfying the criterion of no underlying mutations contributing to the observed epigenetic alterations for these three loci chosen to demonstrate detection of epigenetic differences.

### MAPit-FENGC with ∼ 1-kb reads delineates regulatory relationships between multiple epigenetic modules

Longer reads allow richer examination of epigenetic landscapes, e.g., functional crosstalk between individual regulatory modules at a distance. Therefore, we redesigned 45 of the ∼ 450-nt promoter targets to capture more upstream sequence (range, 930-959 nt; mean, 940 nt; Supplementary Table 1, Sheet 4; and Supplementary Table 2, Sheet 3). These ∼ 940-nt targets were enriched by FENGC of 400 ng and 800 ng from the same two stocks of gDNA from M.CviPI-treated GBM Nx18-25 used to assay the ∼ 450-nt targets. Hi-Fi CCS reads were predominantly the expected ∼ 990 nt (data not shown), of which 77-81% were on target and 38 (84%) targets had ≥ 1 read in each of the 4 libraries (Supplementary Fig. 10a; Supplementary Table 8). Despite the closely matched target lengths, coverage was variable and only 31% (14/45) of the targets retrieved ≥ 26 reads from the combined 800-ng samples (Supplementary Table 9). Nevertheless, H5mCG and G5mCH levels of the 14 targets were highly correlated between the two biological replicates for the same gDNA amount (Supplementary Fig. 10b; *r* ≥ 0.97). Correlation coefficients were also excellent between different gDNA amounts (Supplementary Fig. 11a; *r* > 0.95), as well as between the overlapping regions of the ∼940-nt and ∼ 450-nt targets (Supplementary Fig. 11b; *r* > 0.92).

To further demonstrate concordance of chromatin architectures between the different lengths of molecules and, moreover, to derive additional regulatory insights, we aligned single-molecule heatmaps of the three *EPM2AIP1*-*MLH1* amplicons (Fig. 3a-c). The 450-bp *EPM2AIP1* and 438-bp *MLH1* molecules were mostly unmethylated (Fig. 3a) and open (Fig. 3b) in GBM Nx18-25, whereas in GBM L0 they were strongly hypermethylated and closed (Fig. 1d). Moreover, as in NSC (Fig. 1c), a continuum of positions for both +1 nucleosomes and thus NFR lengths was observed in Nx18-25, which were more constrained for the *MLH1* +1 nucleosome (Fig. 3b, right). In addition, both NFRs harbored a TSS and strong sequence-specific binding of TFs 1 or 2. The 937-bp molecules revealed continuous, variable-length NFRs that encompass *EPM2AIP1* TSS, *MLH1* TSSa, TF footprint 1, and additionally detected TF footprint 3 (Fig. 3c, right). Co-occupancy of the three TFs and two +1 nucleosomes was evident on the long *EPM2AIP1*-*MLH1* molecules. However, the continuum of *MLH1* TSSa +1 nucleosome positions was obscured when clustering of the 937-bp molecules was weighted on the *EPM2AIP1* +1 nucleosome (Fig. 3c, right) and *vice versa* (data not shown). This suggests that the two nucleosomes are mobilized in large part independently of each other, an assertion further supported by weak correlations in accessibility between GCH sites in the two +1 nucleosomes/NFRs (Fig. 3d, bounded by yellow-dashed rectangles). In contrast, considering each +1 nucleosome/NFR separately, higher negative and positive correlations were observed, consistent with nucleosome sliding (Fig. 3d, white-dashed squares). Therefore, the 937-bp *EPM2AIP1*-*MLH1* molecules netted an additional TF footprint and novel regulatory insight — dynamic, but independent, mobilization of two +1 nucleosomes.

**Fig. 3.**
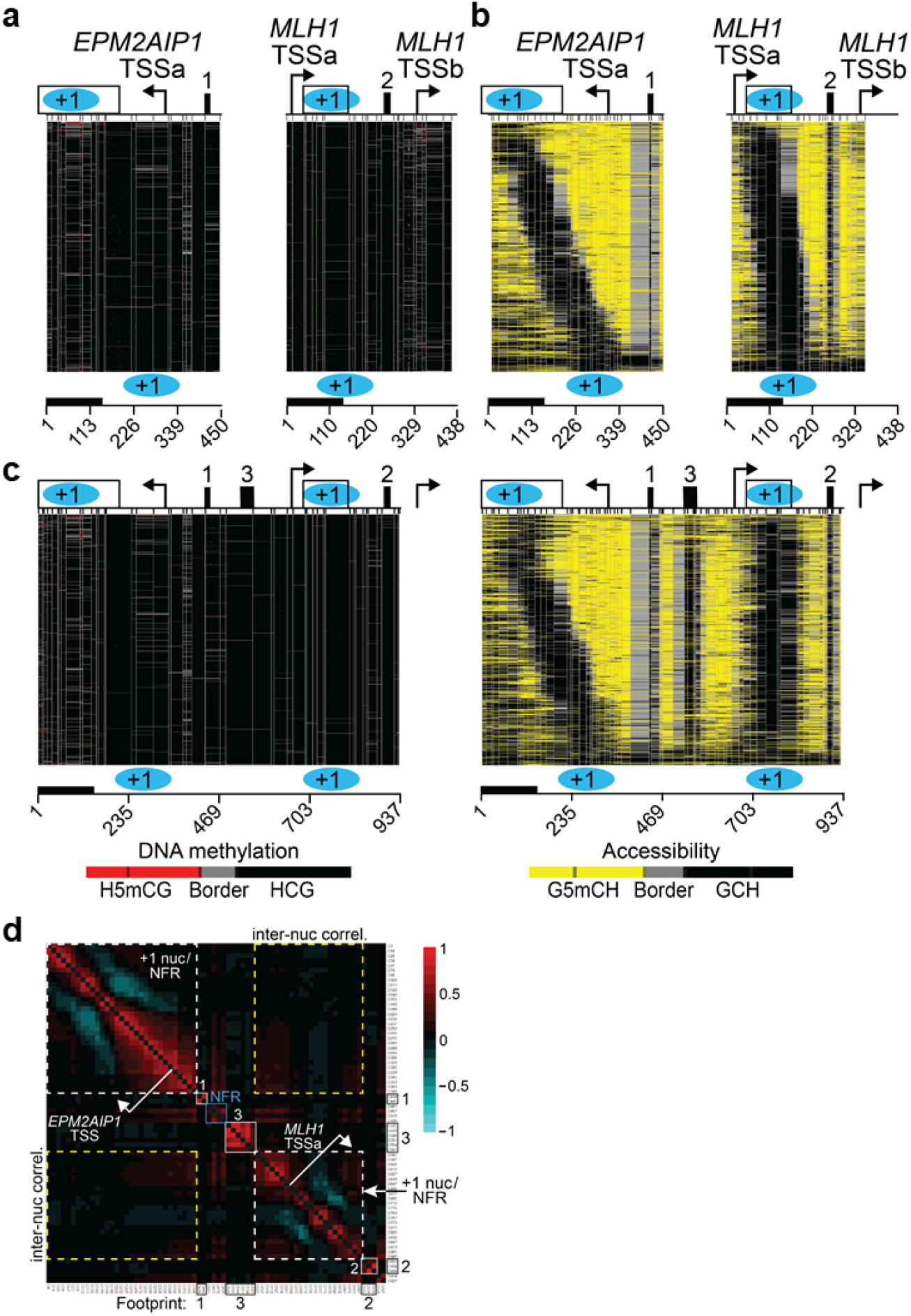
Independent mobilization of the +1 nucleosomes of the divergently transcribed *EPM2AIP1* and *MLH1* promoters. **a-c**, Methylscaper plots of filtered EM-seq CCS reads aligning to *EPM2AIP1* (+307 to **–** 143) (**a**), *MLH1* (**–**13 to +425 relative to *MLH1* TSSa) (**b**), and *EPM2AIP1*-*MLH1* (+328 to **–** 609 relative to *EPM2AIP1* TSS; **–** 650 to +287 relative to *MLH1* TSSa) (**c**). All images contain 1,000 molecules randomly selected from the combined GBM Nx18-25 replicates. The 450-bp *EPM2AIP1* and 438-bp *MLH1* molecules in **a,b** were hierarchically clustered without specifying a subregion and then aligned to the 937-bp *EPM2AIP1*-*MLH1* molecules in **c**. Symbols as defined above. **d**, Correlation of G5mCH levels (each C indicated at bottom and right) of the same 1,000 molecules plotted in **c**. TF footprints in **c** display expected positive correlations (red) adjacent to the diagonal due to co-protection or co-accessibility of adjacent GCH sites (within squares with solid white lines). Off-diagonal, positive correlations within each +1 nucleosome (nuc)/NFR (circumscribed by white-dashed squares) are due to co-protection by histone octamers or co-accessibility in NFRs. Negative correlations (cyan; C70-C155 and C169-C283 in *EPM2AIP1*; C627-C735 and C763-C847 in *MLH1*) arise from inverse protection and accessibility between histones octamers and flanking linkers. Each span of negative correlation corresponds well with the observed range of nucleosome positions at each promoter in **b**. The absence of strong positive or negative inter-nucleosomal correlations (inter-nuc correl.; within yellow-dashed rectangles) supports a conclusion that the two +1 nucleosomes are repositioned independently of each other.

MAPit-FENGC of other ∼ 940-nt targets from GBM Nx18-25 phased additional epigenetic features compared to their shorter counterparts. The divergently transcribed *NPAT* and *ATM* promoters showed a robust TF footprint, a weaker footprint (enclosed by cyan rectangle) at the *NPAT* TSS, and an array of nucleosomes +1 to +3, each progressively less well positioned with increasing distance from the TSS (Extended Data Fig. 3a). The *POLD4* promoter displayed a similar nucleosomal array and two footprints at sequences with strong homology to Sp1 binding sites (Extended Data Fig. 3b, right). Surprisingly, H5mCG localized adjacent to these TF footprints in a sizeable number of molecules (left panel). By contrast to *NPAT*-*ATM* and *POLD4* promoters, the nucleosomes of the *MSH2* 5′ flanking region are more disorganized (Extended Data Fig. 3c). In addition, like *NPAT*, the NFR of a substantial proportion of molecules contains ∼ 55-bp footprints at the TSS (cyan rectangle), consistent with occupancy by RNA polymerase II preinitiation complex^43^. Compared to the shorter *CCN4* promoter molecules from Nx18-25 (Fig. 2f), a subset of longer *CCN4* molecules showed NFRs as long as ∼ 400 bp (Extended Data Fig. 3d); however, intriguingly, no NFRs extended beyond a 66-bp A-rich element. Assaying ∼ 800 bp of 5′ flanking sequence from three of the four genes in Extended Data Figure 3 identified clear transitions between areas of 5mCG depletion and hypermethylation.

### MAPit-FENGC reproducibly informs epigenetic architectures within primary cells

Having successfully applied FENGC to detect epigenetic and genetic variation in cell lines, we tested the epigenetic arm of the protocol on primary, bone marrow-derived monocytes. Cells isolated from four female mice were separately permeabilized and treated with M.CviPI, and 600 ng gDNA from each sample was processed with flap adapters targeting 78 genes with roles in the cellular inflammatory response (Supplementary Table 1, Sheet 5). For this experiment, primers were designed by FENGC oligonucleotide designer (FOLD; Methods). A computational solution for all 78 targets that avoided repeats and satisfied other command-line settings yielded targets ranging from 474-987 nt (mean, 620 nt; Supplementary Table 2, Sheet 4).

Hi-Fi CCS reads from the 4 sequenced libraries were mapped to mm9 build of the mouse genome and the subset of target references: 20-29% did not align to either reference or were removed by filtering, yielding 71-80% on-target reads (Supplementary Table 10). MAPit-FENGC detected 71-75 targets (91-96%) with ≥1 read from each library (Supplementary Tables 10,11).

A mixed effects ANOVA was used to evaluate epigenetic differences between the four monocyte samples (Supplementary Table 12). To provide statistical rigor, testing was restricted to 43 regions with ≥100 filtered CCS reads per sample, of which all were ≤ 760 nt and ≤ 70% GC content (Supplementary Fig. 12). For all 43 targets, no single monocyte sample showed significant differences in DNA methylation or accessibility from the other three samples.

The highly consistent chromatin structures recorded by MAPit-FENGC between individual monocyte samples is apparent in plots of both single-molecules as well as averaged H5mCG and G5mCH for eight representative loci, which also illustrate interesting chromatin biology (Fig. 4; Extended Data Fig. 4). Divergently transcribed *Bop1* and *Hsf1* promoter copies are exceptionally open, with 2,333 of 2,334 total molecules from the 4 monocyte samples displaying relatively large NFRs (range 223-445 bp; Extended Data Fig. 4a). As indicated above, such a locus demonstrates efficient probing of chromatin accessibility with M.CviPI. This supports the observed bimodal distribution of *Btk* molecules (Fig. 4a); epialleles bearing 0% H5mCG and TSS-proximal NFRs with a prominently bound TF, consistent with sustaining transcription *versus* those with variable amounts of H5mCG and reduced, disorganized accessibility reminiscent of X inactivation. Of note, H5mCG is preferentially depleted at the TSS of methylated *Btk* copies that presumably reside on the inactive X chromosome.

**Fig. 4.**
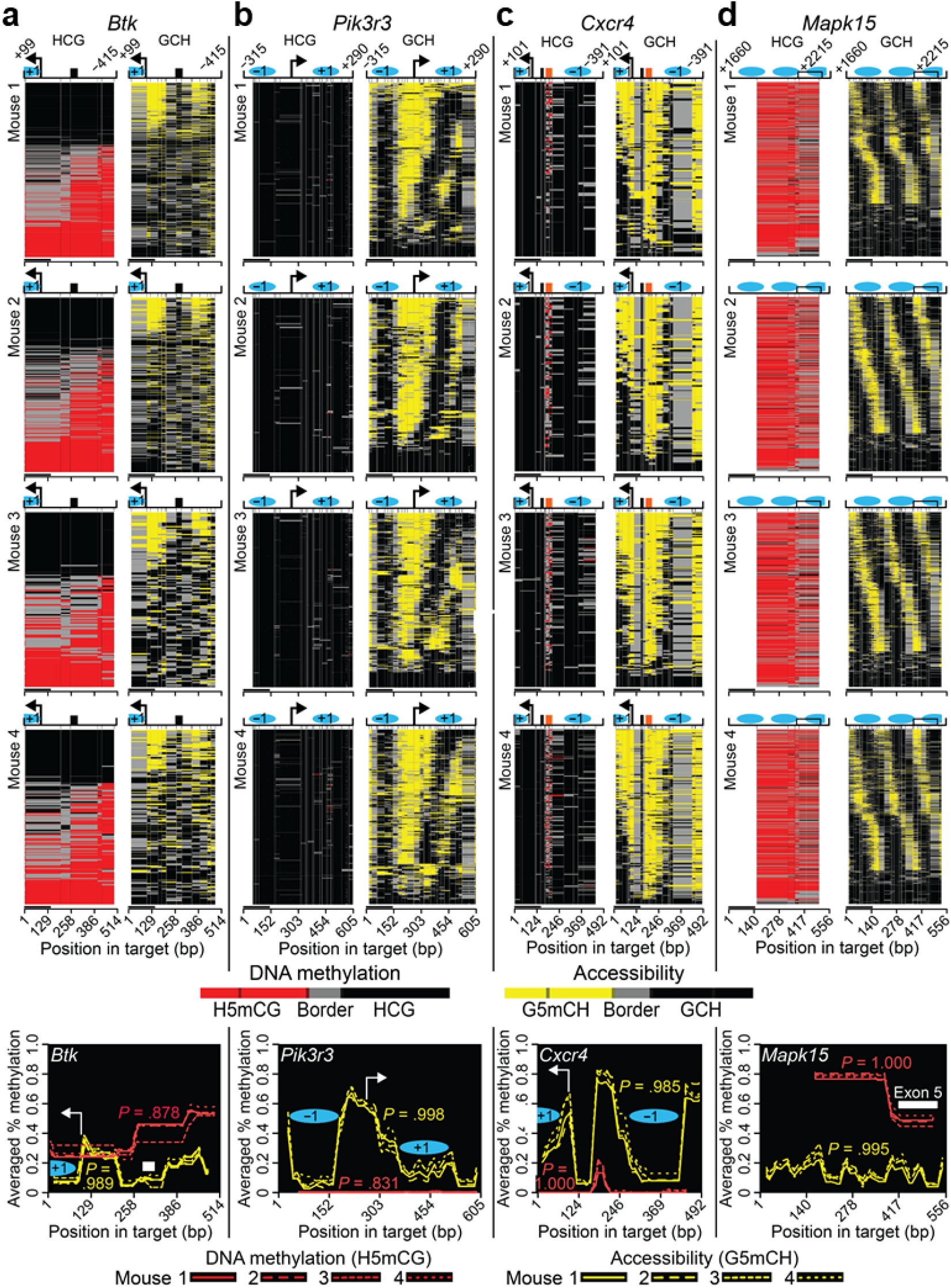
Reproducible, regiospecific chromatin architectures and dynamics within murine bone marrow-derived monocytes. **a-d**, Methylscaper single-molecule methylation heatmaps (top four rows of panels) and averaged levels of H5mCG and G5mCH (bottom row of panels).The plots derive from filtered EM-seq Hi-Fi CCS reads from the sequenced MAPit-FENGC libraries of ∼ 620-nt targets from four female mouse monocyte samples. Single molecules were aligned to the promoters of *Btk* on the chromosome X (**a**), *Pik3r3* (**b**), *Cxcr4* (**c**), as well as the *Mapk15* gene body (**d**). Black rectangles, strong TF footprints; orange rectangle in **c**, 36-bp zone of H5mCG in NFR of *Cxcr4* that is adjacent to but not within the neighboring TF binding site. Base pair position along each target (to scale) is indicated at bottom; black bar on left represents 147 bp. Other symbols as defined above. For these and all other statistically assessed mouse loci, no sample exhibits significant differences in H5mCG or G5mCH from the other three (*P* = 1.000, Bonferroni-corrected, alpha .05; Supplementary Table 12).

Among other intriguing loci, the *Pik3r3* promoter exhibits a well-positioned **–**1 nucleosome and progressive repositioning of the +1 nucleosome, widening or narrowing NFRs in individual cells preferentially on the downstream side of the NFR (Fig. 4b). By comparison, incremental movement of both the **–**1 and +1 nucleosomes produces a ‘tornado-like’ distribution of NFR lengths at the TSSs of *Tlr4* (Extended Data Fig. 4b), *Hsp90ab1* (Extended Data Fig. 4c), *Irf7* (data not shown), and *Cxcr4* (Fig. 4c). Interestingly, the latter promoter displays a 36-bp zone of H5mCG (orange rectangle) in the NFR, located only 12 bp from a strongly footprinted TF.

Compared to the above promoters, *Src* chromatin (**–** 6,420 to **–** 5,805 of RefSeq TSS) is heavily methylated and populated by arrays of randomly positioned nucleosomes with short, accessible linkers (Extended Data Fig. 4d). Interestingly, in the majority of cells, a sequence with strong homology to the consensus CTCF binding site^44^ (map position 462) exhibits a 50-bp area of protection of variable numbers of HCG against methylation. The *Mapk15* gene body (+1,660 to +2,215) is also organized into heavily methylated arrays of randomly positioned nucleosomes, with noticeable depletion of 5mCG in exon 5 (Fig. 4d).

We independently validated MAPit-FENGC of monocytes with single-amplicon MAPit, which reported identical chromatin architectures of the *Btk, Cxcr4, Hsp90ab1*, and *Tlr4* regions (data not shown). Altogether, our data demonstrate that MAPit-FENGC effectively profiles chromatin architectures of purified primary cells, with striking inter-sample reproducibility.

## Discussion

Targeted sequencing offers cost savings, faster turnaround times, and higher sequencing coverages that improve data quality and enable more comprehensive descriptions of specific loci than is feasible with whole genome analysis. However, currently available technologies continue to suffer from limitations regarding target sequence design, high off-target sequencing, and/or prohibitive costs. Here, we describe FENGC, a multiplexed, cost-effective, and high-throughput method, wherein FEN activity precisely excises target sequences reconstituted into double flaps (Fig. 1a). Precision targeted cleavage at any desired nucleotide offers outstanding latitude of target sequence design, mitigates off-target sequencing, and enables length-matching that minimizes amplification and sequencing biases.

Several additional features of FENGC make it particularly attractive for targeted sequencing. First, the reaction is scalable and low cost because it employs standard, salt-free synthesis of target-specific oligos, without modifications such as 5mC or biotinylation. Second, high on-target specificities are achieved by several means (Fig. 1a): i) a pair of sequence-specific flap adapters direct double flap formation, target strand cleavage, and 5’-end ligation; ii) a nested, sequence-specific adapter directs 3’-end ligation; iii) neither flap cutting nor ligation tolerate DNA mismatches^15,17,45-47^; and iv) sequence complexity is considerably reduced by exonuclease digestion prior to amplification, as in nested patch, bisulfite patch, and MAPit-patch PCRs^13,14,48^. Third, for every FENGC target, oligo U1-N within flap adapter 2 ligates to the 5’ end of each downstream sequence. Therefore, a series of flap adapters annealing successively farther downstream and/or upstream enable chromosome ‘walking.’ Fourth, sample loss is minimized through sequential reagent addition, poising FENGC for robotic automation. Therefore, superior sensitivity and on-target frequencies are reproducibly attained at a fraction of the cost of methods employing CRISPR/Cas or pull downs with biotinylated baits.

FENGC-enriched targets can be amplified for genotyping, after C to U conversion for methylation analysis, or both. Conducting genotyping and epigenetic assays in parallel detects mutations that are potential confounders of *bona fide* epigenetic alterations. To this end, sequencing of 119 FENGC-enriched, but non-deaminated targets, from NSC and GBM Nx18-25 treated with M.CviPI detected 43 SNPs and 16 indels already recorded in dbSNP (Supplementary Table 7). This included rs16260, a known *CDH1* SNP and cancer-specific risk factor that impairs transcription^40-42^, as well as eight novel SNPs and one indel.

Non-destructive enzymatic conversion^23,49^ of the same promoter sequences from M.CviPI-treated chromatin, termed MAPit-FENGC, provides integrated, single-molecule detection of DNA methylation and chromatin accessibility, with superior sensitivity to bisulfite conversion (Supplementary Figs. 4b,6). Among promoters meeting stringent CCS read filtering criteria in both bioreplicates of NSC and classical GBM Nx18-25, 24% (13/54) of promoters of genes encoding DNA repair or cancer-associated products exhibited ≥0.1 differential in H5mCG, G5mCH, or both (Supplementary Table 6). Of these 13 promoters, 3 (*GSTA1, HOTAIR*, and *HOXB2*) of 4 with the highest epigenetic differences also possessed 1-3 genetic variants with higher abundances in either non-cancerous NSC or GBM Nx18-25. Such mutations could contribute to observed epigenetic reconfigurations, and their presence should be considered in epigenetic investigations of samples with high mutation burdens.

MAPit-FENGC provides a rich contextual and mechanistic view of epigenetic landscapes by integrating the spatial relationships between endogenous CG methylation, nucleosome positioning/occupancy, and TF binding along contiguous single chromatin fibers. The high coverages provide statistical power for detection of epigenetic differences with a sensitivity as low as 0.1% hypermethylated epialleles (Fig. 1e). Furthermore, hierarchical organization of 100s to 1000s of molecules, each reporting a static snapshot from a single cell, captures the range of epigenetic intermediates and therefore DNA methylation and nucleosome dynamics akin to a flipbook animation. For example, within hypermethylated chromatin, MAPit-FENGC revealed regularly spaced arrays of nucleosomes, which were mobile between different cells, i.e., not positioned relative to DNA sequence (Fig. 4d; Extended Data Fig. 4d). Similar observations were made using single-cell micrococcal nuclease sequencing (scMNase-seq)^50^, albeit MAPit-FENGC expends far less financial and computational resources.

At transcribed mammalian promoters, MAPit-FENGC efficiently captured heterogeneously sized NFRs, as did scMNase-seq using meta-data that average across many loci and cells^50^. However, the superior coverages of MAPit-FENGC also detected *promoter-specific* patterns of incremental nucleosome repositioning and hence changes in NFR length, as we previously reported with MAPit of a synthetic yeast locus bearing various TF binding sites^51^. Such promoter-specific architectures reflect a net outcome of the competitive interplay between nucleosomes and chromatin-associated proteins, including TFs, recruited chromatin remodelers, etc.^51,52^. A complex regulatory interplay seems likely, for example, at the divergently transcribed *EPM2AIP1*-*MLH1* locus, where multiple TSSs, TFs, and mobile nucleosomes reside (Fig. 3). Indeed, long-read MAPit-FENGC divulged that the two *EPM2AIP1*-*MLH1* +1 nucleosomes slide independently and to different extents, challenging the prevailing model that that two genes are coordinately controlled by a shared promoter.

The increased mechanistic granularity of MAPit-FENGC cannot be gleaned from short reads and/or methods that sever the DNA backbone connecting multiple nucleosomes and regulatory elements, including DNase-, MNase-, and ATAC-seq. Single-cell versions of these techniques at comparable coverages of specific loci achieved with MAPit-FENGC are currently not feasible due to low numbers of recovered unique fragments per cell. Fiber-seq^53^, which uses a non-specific N6-adenine methyltransferase to increase footprinting resolution, also has limitations in that simultaneous detection of N6mA and endogenous 5mC is not presently possible. In sum, FENGC enrichment by itself or in combination with MAPit are expected to have broad applications in basic research, biomarker development, plant breeding, etc.

## Data availability

Raw data are deposited in NCBI BioProject under accession PRJNA752452PRJNA752452PRJNA752452.

## Code availability

Code will be made available at https://github.com/uf-icbr-bioinformatics/FOLD.

## Methods

### Cell culture

The human fetal telencephalic NSC line was obtained from Leo A. Behie, who obtained parental consent of donated fetal tissue at the University of Calgary. The use of NSC as a non-cancerous control, as well as establishment of and experimentation with the Nx18-25 cells, was approved by the University of Florida Institutional Review Board 01 Committee (IRB Protocols 201300714, 201500610, and 202002438). NSC and human GBM cell lines (L0 and Nx18-25) were cultured in complete NSC medium (basal medium + proliferation supplement at a 9:1 ratio; NeuroCult™ NS-A Proliferation Kit, STEMCELL Technologies, 05751) supplemented with penicillin-streptomycin (1% final concentration; ThermoFisher Scientific Gibco, 15140122), 20 ng/ml human recombinant epidermal growth factor (STEMCELL Technologies, 78006.1), 10 ng/ml human recombinant basic fibroblast growth factor (STEMCELL Technologies, 78003.1), and 0.679 U/ml heparin (Sigma, H3149)^22^. For NSC culture, 10 ng/ml leukemia inhibitory factor (Millipore, LIF1050) was also added. The cells were maintained in a humidified incubator at 37°C and 5% CO_2_. A standard protocol was used for passaging the NSC^54^ and GBM cells^55^, whereby the neurospheres were collected by centrifugation at 110*g* for 5 min every 7-10 days. The pellet was resuspended in 0.05% (w/v) trypsin, 0.53 mM ethylenediaminetetraacetic acid (EDTA; ThermoFisher Scientific, 25300062), prewarmed to 37 °C. Soybean trypsin inhibitor (ThermoFisher Scientific, 17075029) was then added and gentle pipetting used to dissociate the neurospheres into single cells for replating.

### Animal experiments

Animal protocols were approved by the University of Florida Institutional Animal Care and Use Committee (IACUC Protocols 201807422 and 201910745). ‘Guide for the Care and Use of Laboratory Animals’ was adhered to as prepared by the Committee for the Update of the Guide for the Care and Use of Laboratory Animals of the Institute for Laboratory Animal Research, National Research Council. Mice were closely monitored for signs of dehydration, weight loss, impaired mobility or physiological signs of underlying disorders such as labored breathing or respiratory distress.

Four C57BL/6J female mice were used in order to evaluate detection of distinct transcriptionally active and inactivated epialleles from the X chromosome. The mice were 8-10 weeks of age upon arrival and individually housed on a 12 h dark/12 h light cycle at 19-22°C and 30-60% humidity, with standard chow diet and water provided *ad libitum*. Prior to cell collection at 4 mo age, anesthetization of mice was induced and maintained with 5.0% and 1.5% isoflurane USP (Patterson Veterinary Supply, NDC 14 043-704-06), respectively, using an Eagle Eye Model 150 anesthesia machine (Jacksonville, FL, USA). The depth of anesthesia was monitored by the absence of pedal withdrawal reflex.

### Isolation of monocytes from mouse bone marrow

Bone marrow from four mice was collected from spines (below skull to above tail) cleaned of extraneous tissues. All subsequent steps were conducted under sterile conditions. First, the tissue was crushed at room temperature using a ceramic mortar and pestle in a sterile solution of 10 ml of phosphate-buffered saline (PBS), pH 7.2, 2 mM EDTA, 0.5% (w/v) bovine serum albumin (BSA), by mixing MACS BSA Stock Solution (Miltenyi, 130-091-376) and Biotec, autoMACS^®^ Rinsing Solution (Miltenyi, 130-091-222) in a ratio of 1:20. Subsequently, the crushed spine was removed, and the homogenate was filtered through pre-separation filters with 30-μm nylon mesh (Miltenyi Biotec, 130-041-407,) into a 15-ml Falcon^®^ tube. The cellular filtrate was then washed by centrifugation at 300*g* for 10 min, removal of the supernatant, and resuspended in the same PBS/BSA solution for cell counting (Heska, Element HT5 Hematology Analyzer). Monocytes were then enriched using a negative isolation protocol specific to mouse bone marrow (Miltenyi Biotec, 130-100-629), according to the manufacturer’s protocol. Briefly, ∼40-80 million washed cells were incubated with FcR blocking reagent and cocktail of mouse biotinylated-antibodies, washed with PBS/BSA, resuspended in degassed RPMI 1640 medium (ThermoFisher, 11875101) containing 1% (w/v) penicillin-streptomycin (ThermoFisher Gibco, 15140163), and efficiently depleted of non-target cells by addition of magnetic microbeads conjugated to mouse anti-biotin monoclonal IgG1 antibodies and passage through LS ferromagnetic columns (Miltenyi Biotec, 130-042-401). The flow through, containing highly enriched bone marrow-derived monocytes, was collected into a 1-ml centrifuge tube. To allow recovery from the collection process, a mean of 1.2 million monocytes were plated in a well on a 96-well plate and incubated in a humidified 37°C incubator at 5% CO_2_ in RPMI 1640 containing 1% penicillin-streptomycin solution for 3 h before harvesting for MAPit-FENGC.

### FEN cleavage of oligonucleotide targets

Cleavage efficiencies of *Thermococcus* 9° N FEN1 and Taq (Supplementary Fig. 1b) were tested on a 129-nt 5’ flap assembled by incubating 500 nM each of 200-mer T target, 200 oligo 1-T, and oligo U1 (Supplementary Table 1, Sheet 1). The 200-mer T sequence corresponds to a segment of pGEM-3Z/601b^56^ with A 2453 substituted with T by site-directed mutagenesis with oligos TGCACC<u>T</u>TATGCGGTGT and CTCTCAGTACAATCTGCTCTGA. Reactions also contained 1 U APEX Taq (Genesee Scientific), 1 U HotStar Taq (Qiagen; based on polymerization activity), and 32 U FEN1 (New England Biolabs). The buffers used were: 1x PCR buffer (Qiagen; referred to as Taq buffer); 1x ThermoPol reaction buffer (New England Biolabs; referred to as FEN1 buffer); and the mixed buffer reactions contained final concentrations of 1x CutSmart buffer (New England Biolabs) and 1x Taq buffer or FEN1 buffer. The incubation conditions were 3 min at 95°C, followed by 20 min at 65°C and then 9 cycles of 0.5 min at 95°C and 10 min at 65°C. Digestion products were separated on an Agilent 2100 Bioanalyzer and the areas under the peaks of cut and uncut 200-mer T were integrated to determine the cleavage percentages.

To test the effects of different ‘overlapping’ nucleotides and DNA methylation on FEN1 activity, two 80-mer targets of identical sequence were synthesized with and without five 5mC residues at the same locations (Supplementary Fig. 2a; Supplementary Table 1, Sheet 1). These 5 cytosines occupy the upstream duplex contacted by human FEN1 co-crystallized with a 5’ flap and 1-nt 3’ flap^19^. The two 80-mers (50 nM) were assembled in separate 20-μl reactions with their corresponding unmethylated flap adapters (500 nM each of oligo U1-A, C, G, or T and 80 oligo-A, C, G, or T; Supplementary Fig. 2b,c; Supplementary Table 1, Sheet 1). This reconstitutes A, C, G, or T 5’ of the cleavage site and as the 1-nt 3’ flap, as well as recapitulates the scenario of performing FENGC with methylated DNA (Supplementary Fig. 2a). Reactions also contained 1 x FEN1 buffer and 0, 0.01, 0.04, 0.2, or 1 unit(s) of FEN1 and were incubated for 3 min 95°C, followed by 20 min at 65°C. The percentages of cleaved 80-mer at increasing doses of FEN1 were determined by quantitative, real-time PCR, calculated by [1-2^-(Ct of FNE1 – Ct of no FEN1)^] x 100. The dose *versus* response curves (Supplementary Fig. 2b) and EC_50_ values (Supplementary Fig. 2c) were plotted using GraphPad Prism 5 (www.graphpad.com).

### Methylation of accessible GC sites for single-molecule MAPit methylation footprinting

Two million cells were harvested with 0.05% (w/v) trypsin, 0.53 mM EDTA (ThermoFisher Scientific, 25300062) and washed once with ice-cold PBS, pH 7.2, supplemented with 0.015% (w/v) sodium azide. Cells were then centrifuged for 5 min at 1,000*g*, washed once with 500 μl ice-cold cell resuspension buffer (20 mM HEPES, pH 7.5, 70 mM MgCl_2_, 0.25mM EDTA pH 8.0, 0.5mM EGTA pH 8.0, 0.5% (v/v) glycerol, freshly supplemented 10 mM DTT and 0.25 mM PMSF). After centrifugation for 5 min at 1,000*g*, cells were resuspended in 180 μl cell resuspension buffer with supplemented with 0.05% (w/v) digitonin and incubated on ice for 10 min. Cells were stained with trypan blue to ascertain 100% permeabilization. Cells were then divided into two aliquots and treated with and without 100 U M.CviPI GpC methyltransferase (100 U/million cells; New England Biolabs, M0227B-HI) supplemented with fresh 160 μM *S*-adenosyl-*L*-methionine for 15 min at 37ºC. Reactions were stopped by addition of an equal volume of 1% (w/v) SDS, 100 mM NaCl, 10 mM EDTA and vortexed briefly at medium speed. Nuclei were treated with 100 μg/ml RNase A for 30 min at 37ºC followed by 100 μg/ml proteinase K treatment at 50ºC overnight. Genomic DNA was extracted using phenol-chloroform-isoamyl alcohol (25:24:1, v/v) phase separation, followed by ethanol precipitation, and resuspension in molecular-grade H_2_O.

### FENGC

Except as noted otherwise, the human gene panels targeted the promoter TSSs of genes coding for proteins with DNA repair and/or cancer-associated functions. Flap oligos 1 and 2 as well as nested oligos 3 were designed for 11 targets of ∼ 300-nt, 119 targets of ∼ 450-nt, and 45 targets of ∼ 940-nt, as catalogued in the respective sheets of Supplementary Tables 1 and 2. The mouse panel targeted the promoters of 78 inflammatory response genes, unless noted otherwise.

To mitigate 5′ flap secondary structure, purified gDNA from M.CviPI-treated cells was fragmented by sonication with a UCD-200 Bioruptor (Diagenode) at high level for 25 sec in 100 μl molecular-grade H_2_O. Subsequently, the volume was brought to 20 μl in a SpeedVac vacuum concentrator.

Sequence-specific flap oligos 1 and 2 as well as oligo 3 (4 nmole each) were ordered as 0.3-ml each in 96-well format as salt-free oligos (Eurofins). Lyophilized oligos were resuspended at 100 μM concentration in molecular-grade water. Unless indicated differently, 500 ng gDNA was processed in each reaction. Multiplexed cleavage of 5′ flaps was performed by adding 1 μl 10 μM oligo U1-T, 2 μl pooled oligos 1, 2 μl pooled oligos 2, 1 x Qiagen PCR Buffer (Qiagen), 3 U APEX Taq (Genesee Scientific), and molecular-grade H_2_O to bring the total volume to 35 μl.

Reactions were incubated for 3 min at 95°C, then 20 min 65°C, followed by 14 cycles of 0.5 min at 95°C and 10 min at 65°C. Subsequently, 2 μl pooled oligos 3, 1 μl 10 μM oligo U2, 1 x Ampligase® Reaction Buffer (Lucigen), 10 U Ampligase (Lucigen), and molecular-grade H_2_O were added to bring the total volume to 45 μl. Incubation of the reactions at 95°C for 3 min was followed by 30 s at 95°C and 8 min at 65°C for 100 cycles. Unprotected DNA was removed by addition of 20 U exonuclease I, 100 U exonuclease III (both from New England Biolabs), and incubation for 60 min at 37°C and 20 min at 80°C.

The reduced representation products were subjected to either standard PCR, BS-PCR, or EM-PCR. For standard PCR, captured DNA was purified with MinElute PCR purification kit (Qiagen) or AMPure XP beads (1.8x), and PCR was performed with HotStar Taq (Qiagen) as described above. For BS-PCR, captured sequences were converted with EZ DNA Methylation Gold Bisulfite Treatment Kit (Zymo Research) as indicated above, except that 35 cycles of amplification were used. For EM-PCR, captured sequences were purified with 1.8x AMPure XP beads (Beckman Coulter) or 1.8x NEBNext beads (New England Biolabs). Enzymatic conversion of cleaned sequences was done with NEBNext^®^ Enzymatic Methyl-seq Conversion Module (New England Biolabs, E7125), according to the manufacturer’s instructions, and converted sequences were eluted in 14 μl molecular-grade H_2_O. PCR was performed the same way as BS-PCR. For EM-PCR of ∼ 940-nt targets, sequences were eluted with 25 μl molecular-grade H_2_O. PCR was performed with 500 nM of oligo U1 and U2 primer in 2x KAPA HiFi HotStart Uracil+ ReadyMix (Roche) in 50 μl. Amplification conditions for BS- and EM-PCR were 5 min at 95°C, followed by 30 cycles of 20 sec at 98°C, 15 sec at 62°C, 30 sec at 72°C, and a final extension of 1 min at 72°C. Amplified FENGC products were purified with AMPure XP beads (0.65x for ∼ 500-bp products; 0.75x for ∼ 350-bp products).

To validate FENGC of gDNA from M.CviPI-treated cellular chromatin, i.e., MAPit-FENGC, single-amplicon MAPit was performed as previously described^20^, except EM-PCR was used as indicated above. In a MAPit-FENGC reaction, all enriched sequences are used for EM-PCR. Therefore, an aliquot of the same gDNA purified from the four M.CviPI-probed mouse monocyte samples was freshly enzymatically converted and the indicated primer pairs (converted bases in lower case) were used to amplify the promoters of *Btk* (+140 to − 451; ttAAAtttAttTtAGTTtTGAtTTAAT and aTCTCTaAaTaArCCTTCTTTaTCTa), *Cxcr4* (+132 to − 418; GTAGGATGyAAGTGGAtTTAtAtTtAt and aaAAaAaTTTTaaTAaTAATCAC-TCCTaAC), *Hsp90ab1* (+252 to − 401; GTAtAGyttGAAttTtTAAttT and AaTTarCCACCTCC-ATTCTCT), and *Tlr4* (− 248 to +379; ATtATGAtAtAAGAtAtGGyAAtT and ACACATACCTC-TATrCAaaaATTCAA). These regions are nearby but not the same as those analyzed by MAPit-FENGC (Fig. 4; Extended Data Fig. 4), in order to avoid as many CG and GC as possible. FENGC with BS- or EM-PCR can ignore DNA methylation status, highlighting another advantage of the method.

### Pacific Biosciences Sequel and Sequel IIe SMRT sequencing

Purified amplicons passing quality evaluation on the Agilent TapeStation high-sensitivity DNA5000 tape were submitted to the University of Florida Interdisciplinary Center for Biotechnology Research (UF-ICBR, RRID:SCR_019152) for SMRT bell library construction (P/N 101-791-700 Version 07). The manufacturer’s protocol was used, except the input amount of each amplicon was increased 3-6x for DNA damage repair, end repair, A-tailing, and barcode ligation. Equimolar amounts of barcoded amplicons were pooled and ligated to SMRTbell adaptors, which were then pooled in equimolar amounts and treated with exonuclease, using the SMRTbell Enzyme Clean-Up Mix (PacBio, 101-932-600). Library construction typically yielded sufficient SMRT bell library (∼ 70 ng) for 2-3 SMRT cells.

Sequencing on the PacBio Sequel or Sequel IIe was performed by UF-ICBR and on the Sequel IIe at the Georgia Genomics and Bioinformatics Core at the University of Georgia. The library pool was loaded at 120 pM, using diffusion loading and 20-to 30-h movies, with HiFi generation and demultiplexing. Sequencing Kit 2.0 (PacBio, 101-389-001) and Instrument Chemistry Bundle Version 11.0 were used. All other steps were performed according the recommended protocol by the PacBio sequencing calculator. One Sequel IIe SMRT cell typically resulted in 3-4 million polymerase reads.

High-fidelity circular consensus sequencing (CCS) reads were generated at ≥ 3 single polymerase read passes to determine on-target read efficiencies, with other default parameters. To do so, CCS reads were aligned to the human genome (build GRCh38) using the BWA aligner (bwa-mem)^57^. For reads from deaminated samples, the alignment was performed using bwameth^58^, which uses the BWA aligner. Reads that aligned anywhere to the human genome were then extracted and aligned to the target references, again using either BWA or bwa-meth. On-target efficiencies were then calculated as the percent of aligned reads. Both BWA and bwa-meth were used with default parameters.

For epigenetic analyses, CCS reads were generated with default parameters, except for a setting of ≥ 5 single polymerase read passes. CCS reads were aligned to reference sequences with the python reAminator pipeline^59^. Cut-offs of 95% conversion rate and 95% length of reference sequences alignment were applied. To distinguish endogenous CG methylation and M.CviPI-probed GC methylation unambiguously GCG sites were omitted for calculation of methylation levels, i.e., HCG and GCH, where H is A, C, or T. Single-molecule heatmaps of HCG and GCH methylation were hierarchically clustered (unsupervised) and plotted using methylscaper^26^.

For genotyping, CCS reads were obtained with at least 3 passes and minimum predicted accuracy of 0.99. Reads alignment was performed using minimap2 and SNPs and indels calling were carried out using GATK4 pipeline^60^. All variants were filtered with thresholds of GQ> 30 and Depth> 20, as well as AS_QD< 2.0 for SNPs and AS_QD< 5.0 for indels.

### Reverse transcriptase quantitative PCR (RT-qPCR)

RNA was extracted from one million cells using Direct-zol RNA Miniprep Kit (Zymo Research, 11331). Complementary DNA was synthesized using SuperScript™ First-Strand Synthesis system (Thermo Fisher Scientific, 11904018), then diluted 1:1 with molecular-grade H_2_O and used for qPCR measurement. QPCR was done using Taqman assay (Life Technologies 44-449-63) with gene-specific probes *CCN4* (*WISP1*; Thermo Fisher Scientific Hs00180245_m1), *CD44* (Thermo Fisher Scientific Hs01075861_m1), and *HIST1H1B* (Thermo Fisher Scientific Hs01075861_m1).

### FENGC Oligonucleotide Designer (FOLD)

For FENGC of the ∼ 620-nt target panel (Supplementary Table S2, Sheet 4) from mouse monocytes, the primers (Supplementary Table S1, Sheet 5) were designed by newly developed program, FOLD (https://github.com/uf-icbr-bioinformatics/FOLD). The programs searches an input file of gene names or genome coordinates for primers that avoid repeats and satisfy criteria of Figure 1, such as locating the ‘overlapping’ residues that create 1-nt 3′ flaps. Other command-line options include, but are not limited to, increasing the length of default 500-nt sequences and percentage tolerance of departure from this specified, specification of annotated TSS (e.g., RefSeq), and minimum and maximum primer melting temperature (T_m_).

FOLD implements a sophisticated optimization method to find the best combination of primers for each input sequence (local optimization), and for all input sequences at the same time (global optimization). For each input sequence, the program identifies all possible locations for each one of the three primers, and finds the optimal triple, taking into account the predicted T_m_ for each primer and the length of the corresponding products. The deviation from ideal T_m_ and ideal product length are combined through user-defined weights, and the set of primers with the smallest resulting score is selected.

The goal of global optimization is to ensure that the sizes of the produced fragments for all target sequences are as similar as possible, i.e., exhibit the smallest variance. This is accomplished by picking a random set of primers for each target (among the ones that fulfill the specified design constraints), computing the variance of the resulting fragment sizes, and repeating the process a large number of times, and finally picking the combination that gave the smallest overall variance. Sub-optimal primer sets are therefore accepted for some targets in order to attain more uniform fragment sizes at the global level.

The program’s main output is a table listing the coordinates and size of the resulting PCR product, and information on the three primers (including sequence, length, T_m_, and GC%) for each target in input. Optionally, the program can also write the sequences of all PCR products and of all primers to user-specified files in FASTA format.

### Statistics and Data Analysis

To evaluate the 0.1% NSC gDNA:99.9% GBM L0 gDNA mixture (Fig. 1e), a proportions test was conducted in R using the command prop.test(), with a null hypothesis that the observed proportion of 10 hypermethylated out of 1,781 unmethylated epialleles is less than 0.1%. The *P* value of < .0001 indicates that the proportion of hypermethylated epialleles detected was at least or greater than 0.1%.

To identify differentials in H5mCG (methylation) and G5mCH (accessibility) between NSC *versus* GBM Nx18-25, generalized estimating equations were used to model the effect of cell line on the proportion of methylation per molecule. Only targets with ≥ 50 total CCS reads in the combined replicates (≥ 22 in any single replicate) were considered (Supplementary Table 5). Errors were modeled as normally distributed, and the correlation structure was assumed exchangeable within each replicate. The geeglm function from the geepack v1.3-2 R package^61^ was used in R version 4.1.0. *P* values were corrected for multiple testing using the Bonferroni method to control the family-wise error rate.

To test if MAPit-FENGC detected significant epigenetic differences in at least one of the four analyzed mouse bone marrow-derived monocyte samples, smoothed moving averages (20-bp window) of DNA methylation and accessibility across each gene region were modeled using a mixed effects ANOVA^62^. Testing was restricted to 43 amplicons with ≥ 100 filtered CCS reads per sample and good diversity, i.e., absence of many duplicates (Supplementary Table 10). The mixed effects ANOVA model was fit using the gls function in the nlme v3.1-152 package in R version 4.1.0. Each sample was treated as a random effect, and correlation along the gene region was modeled as that from an autoregression moving average model (ARMA). Autocorrelation parameters were estimated using the auto.arima function from the forecast v8.15 R package, with a maximum possible value of 1 to avoid overfitting. If the differencing order was estimated as zero, then first-order differences were taken of DNA methylation and accessibility. If the autoregressive and moving average parameters were both estimated as zero, then an autoregressive model of order one was used. The *P* values are of the interaction term of base pair and replicate, which tested for differences between replicates across the gene region. *P* values were corrected for multiple testing using the Bonferroni method to control the false discovery rate (Supplementary Table 10).

The violin density plots were made using the “yarrr” R package^63^. The median (horizontal line), interquartile range of methylation levels (box), and smoothed probability density at different methylation levels (gray area) are plotted. *P* values are corrected for multiple testing using the Bonferroni method (α = 0.05) to control the false discovery rate (Supplementary Table 6).

## Acknowledgements

We thank Leo Behie for generously providing the NSC. We are grateful to the Next-Generation DNA Sequencing (RRID:SCR_019152) as well as Gene Expression and Bioinformatics Cores RRID:SCR_019145) of the Interdisciplinary Center for Biotechnology Research of University of Florida for their technical support. This work was supported by grants HDTRA1-16-1-0048 awarded by the Defense Threat Reduction Agency to P.C. and R01 CA155390 awarded by The National Institutes of Health to M.P.K. We also thank the National Brain Tumor Society and the Florida Center for Brain Tumor Research for their support.

## Author information

## Contributions

N.H.N. and M.P.K. conceived and designed the FENGC approach, and N.H.N. conducted initial proof-of-concept experiments with plasmids. M.Z. and M.P.K. conceived the study and designed experiments that M.Z. conducted to develop the mature FENGC protocol. H.Z. and B.A.R. established the novel GBM cell lines. K.O.M. and T.L.C. prepared the primary mouse monocytes and selected the inflammatory response gene targets. A.W. and M.-P.L.G. performed MAPit and qPCR. J.O.B., A.R., J.R.B.N., and W.S.O. provided bioinformatics and computational support, as well as data analyses. A.R. also wrote program FOLD. R.B. conducted biostatistical analyses. F.J.P.-P. and A.C. analyzed the genotyping sequencing results and prepared the data table. M.Z., P.C., and M.P.K. wrote the manuscript. All authors reviewed, edited, and approved the manuscript.

## Inclusion and Ethics

The research included local researchers throughout the research process, whose roles and responsibilities were agreed to amongst the collaborators in advance of conducting the research. During the course of the work, every effort was made to mentor colleagues and promote their career advancement. Local and regional research relevant to our study was taken into account in the citations.

## Competing interests

M.Z., N.H.N., and M.P.K. are co-inventors (assignee, University of Florida) of pending patent application (PCT/US2022/020624, filed March 16, 2022) for the FENGC technology.

## EXTENDED DATA FIGURES

**Extended Data Fig. 1.**
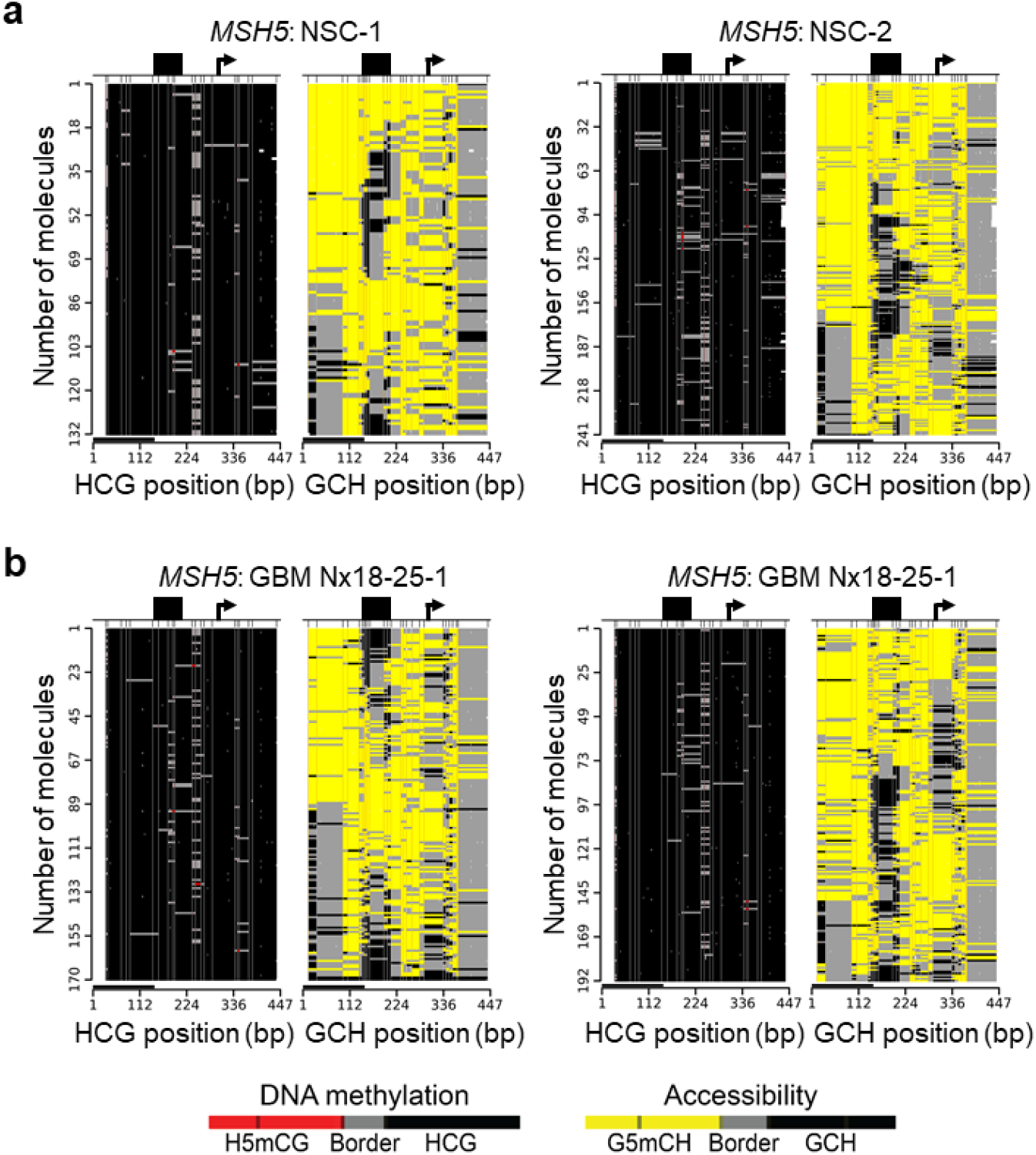
*MSH5* promoter exemplifies a markedly accessible chromatin that controls for inter-sample and -molecule M.CviPI probing efficiency. **a,b**, Methylscaper plots of filtered CCS reads aligned to the *MSH5* promoter (**–** 302 to +145) from MAPit-FENGC with the 119-target panel of two independent cultures of human non-cancerous NSC (132 and 241 reads; Supplementary Table 5) (**a**) and GBM Nx18-25 (170 and 192 reads) (**b**). Each HCG and GCH position (in base pairs) along the *MSH5* amplicon is indicated at the bottom of its respective panel (black bar represents 147 bp). A heterogeneous-sized footprint is marked by a black rectangle. Almost all *MSH5* promoter molecules across all 4 samples (733 of 735) had ≥10 methylated GCH sites, demonstrating high-level cell permeabilization and M.CviPI activity.

**Extended Data Fig. 2.**
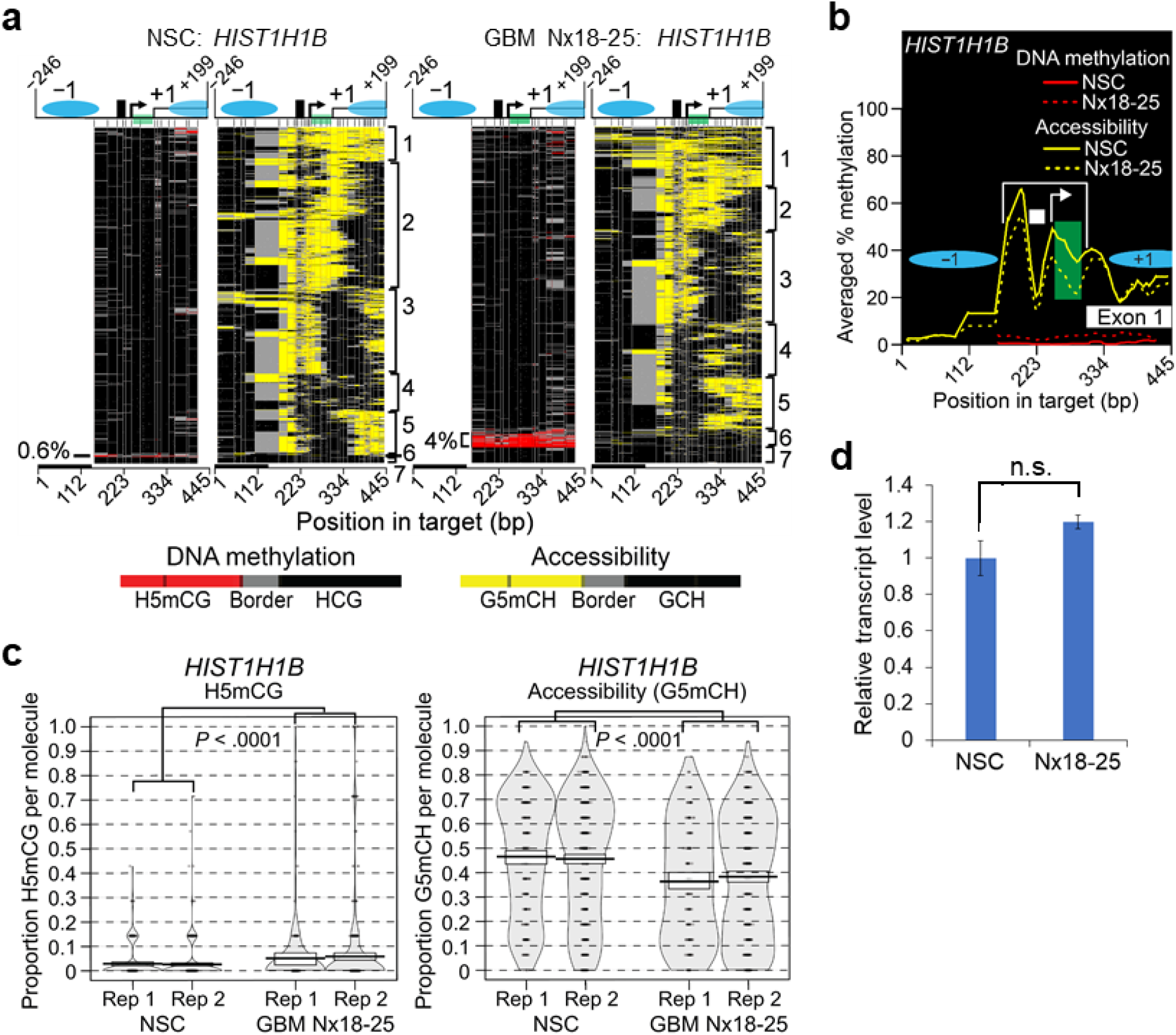
MAPit-FENGC with EM-seq detects minority subpopulations of molecules with differential epigenetic configurations. **a-c**, Methylscaper single-molecule heatmaps (**a**), percentages of averaged H5mCG and G5mCH (**b**), and violins plots (**c**) derive from filtered CCS reads aligning to *HIST1H1B* from two human NSC (*n* = 2; 926 combined reads; Supplementary Table 5) and two GBM Nx18-25 cultures (*n* = 2; 953 combined reads). The bracketed area from **–** 80 to +45 in **b** shows a significant decrease in accessibility in Nx18-25 relative to NSC that meets our 0.05 threshold (**–** 0.074; Supplementary Table 6; *P* < 0.0001). **d**, Relative expression of *HIST1H1B* (*n* = 3, mean ± SD; n.s., not significant). The ∼30-bp footprint upstream of the TSS (black rectangle) is consistent with high occupancy of this consensus TATA box by the preinitiation complex ^43^. The downstream ∼ 50-bp footprint (green rectangle) may correspond to paused RNA polymerase II^43^. Other symbols as defined above.

**Extended Data Fig. 3.**
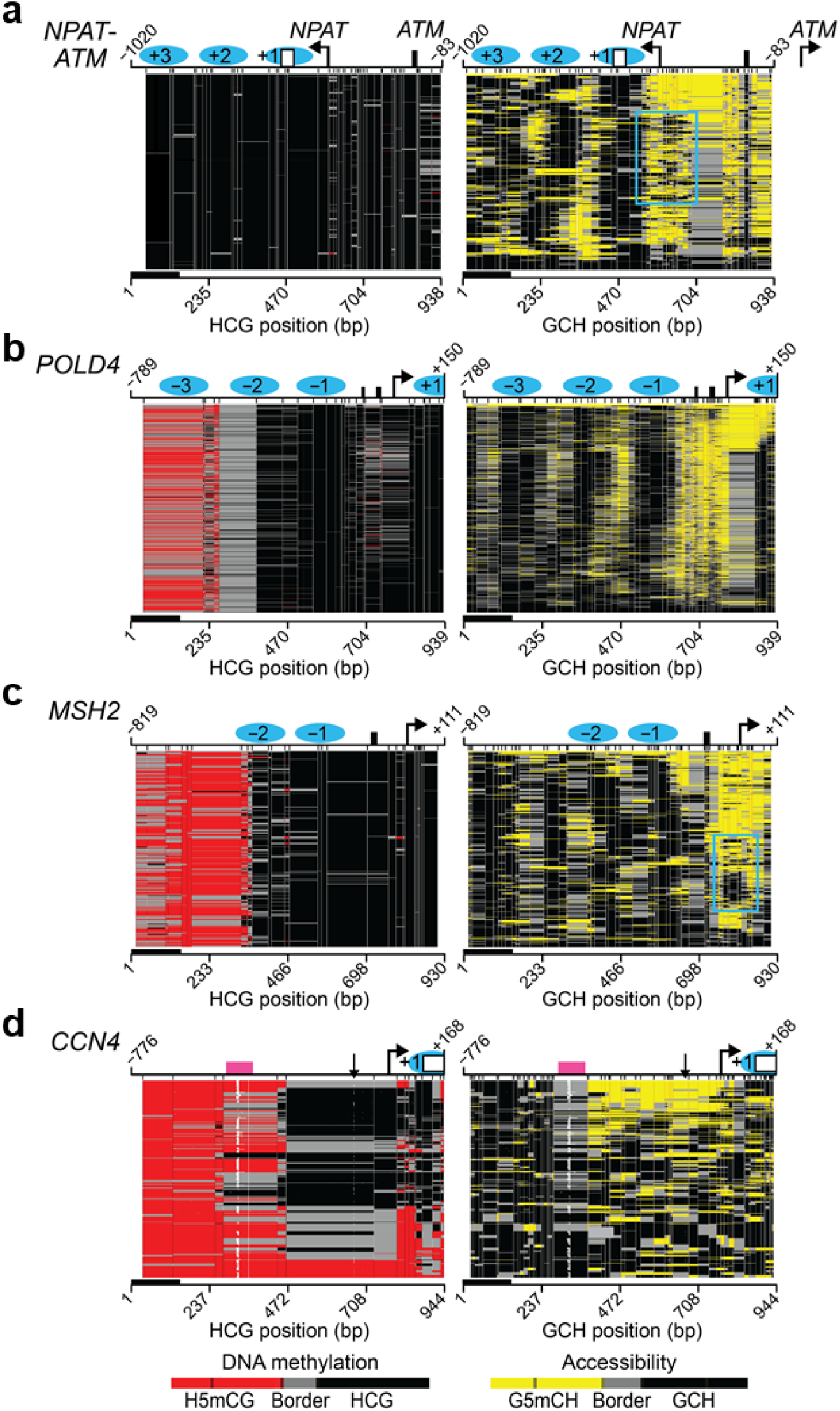
Long-read MAPit-FENGC phases multiple epigenetic features. **a-d**, Methylscaper heatmaps of filtered EM-seq CCS reads aligning to *NPAT*-*ATM* (**–** 1020 to **–** 83 relative to *ATM* TSS; +592 to **–** 346 relative to *NPAT* TSS) (**a**), *POL4D* (**b**), *MSH2* (**c**), and *CCN4* (**d**). All plots contain 1,000 molecules randomly selected from the combined GBM Nx18-25 replicates. Hierarchical clustering was focused to display the longest NFRs. Relatively heterogeneous footprints (cyan rectangles) co-localizing with the *NPAT* TSS in the GCH panels in **a** and **c** are consistent with occupancy by the RNA polymerase II preinitiation complex^43^. In **d**, a vertical arrow points to a known GA to G indel (dbSNP, rs548251181; Supplementary Table 7) that is plotted in white due to local non-alignment with the hg38 reference sequence. Similarly, in **d**, variable-length spans of non-alignment to a 66-bp A-rich sequence (88% A; pink rectangle) are plotted in white. Bent arrows, TSSs; cyan ovals, nucleosomes; black rectangles, TF footprints; white rectangles, protein coding sequences.

**Extended Data Fig. 4.**
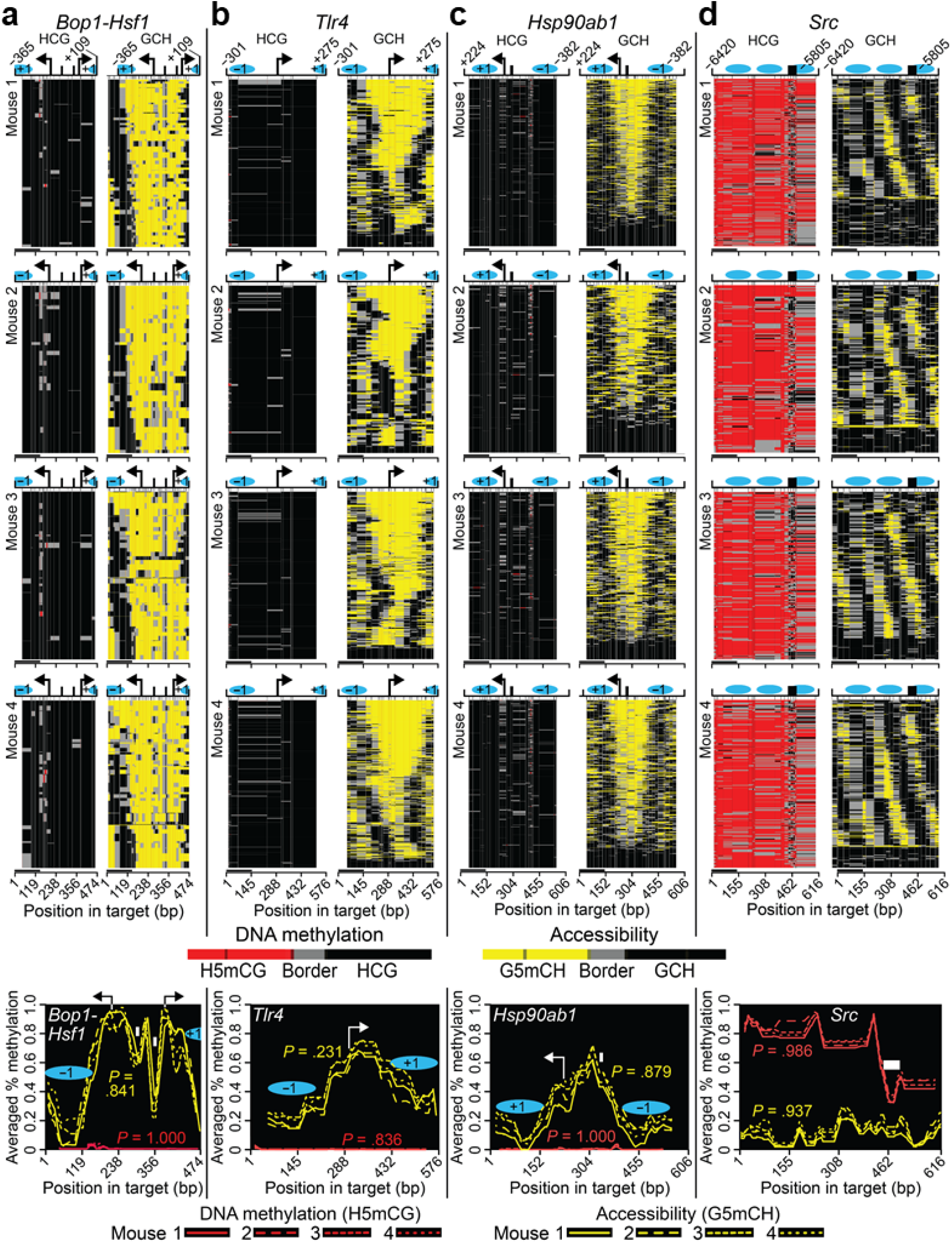
Reproducible, regiospecific chromatin architectures and dynamics revealed by long reads. **a-d**, Methylscaper single-molecule methylation heatmaps (top four rows of panels) and averaged levels of H5mCG and G5mCH (bottom row of panels). The plots derive from filtered EM-seq Hi-Fi CCS reads from the sequenced MAPit-FENGC libraries of ∼ 620-nt targets from four female mouse monocyte samples. Single molecules were aligned to the promoters of divergently transcribed *Bop1* and *Hsf1* (**a**), *Tlr4* (**b**), *Hsp90ab1* (**c**), as well as **–** 6,420 to **–** 5,805 relative to the *Src* RefSeq TSS (**d**). Distances in **a** are relative to the *Hsf1* TSS at right. Black rectangle in **d**, H5mCG footprint with strong CTCF binding site homology^44^. Position along each target (to scale) is indicated at bottom; black bar on left, 147 bp. Other symbols as defined above. For these and all other mouse loci, no sample is significantly different in H5mCG or G5mCH from the other three (*P* = 1.000, Bonferroni-corrected; Supplementary Table 12).

## SUPPLEMENTARY FIGURES

**Supplementary Fig. 1.**
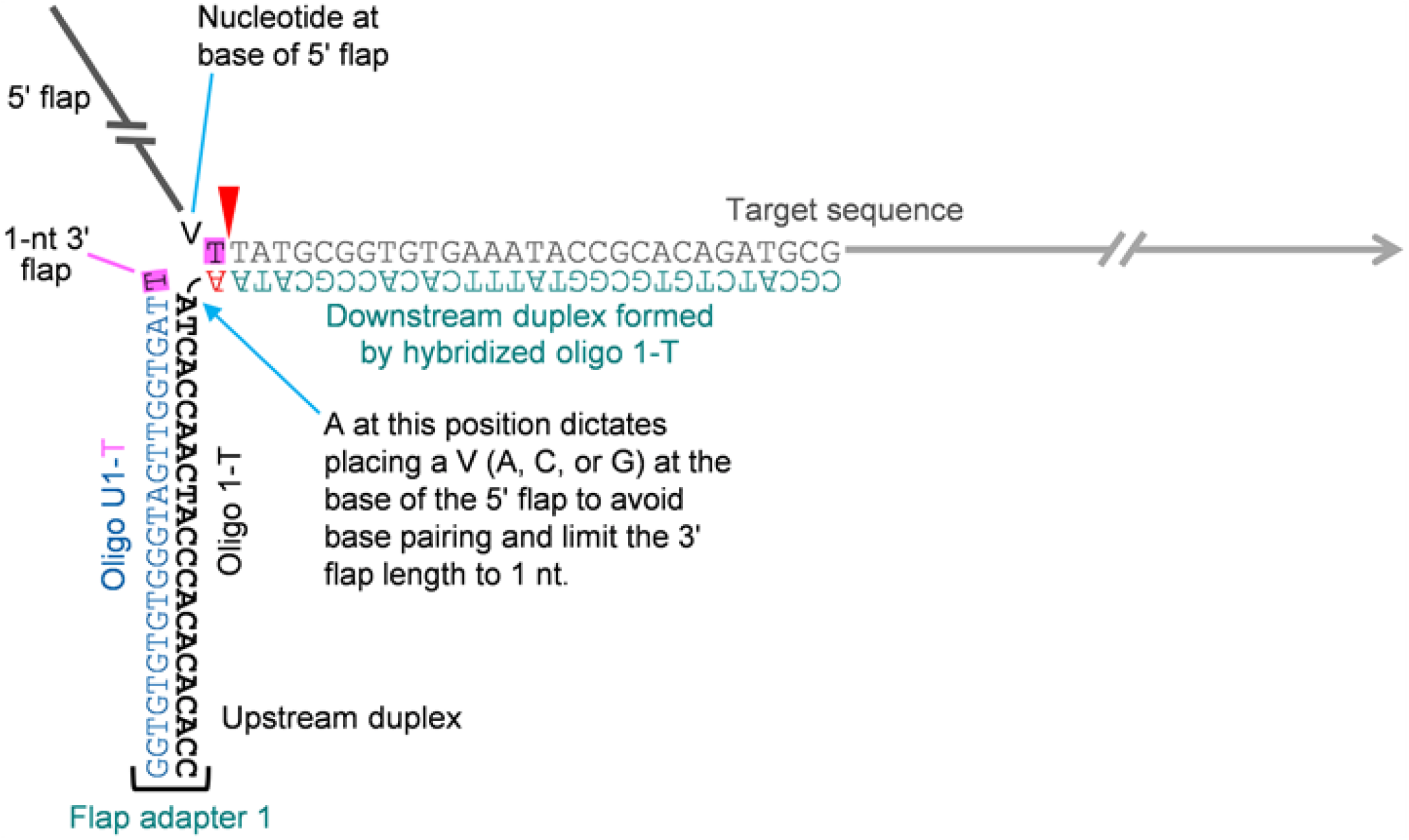
Flap adapter structure to direct efficient, precise 5’ flap scission and target strand release. Shown is part of 200 oligo 1-T hybridized to the 5’ end of its respective target sequence (200-T oligo, gray), forming a downstream duplex. In multiplexed FENGC, an oligo 2-T also hybridizes to the 3’ end of each target sequence (not shown). The constant 3’ tail of all oligos 1-T and 2-T binds to oligo U1-T, forming an upstream duplex and reconstituting a plurality of double flaps. The ‘overlapped’ nucleotides (highlighted pink) of the oligo U1-N 3’-terminus and immediately 5’ of the scissile phosphodiester contain the same base. In the shown configuration using oligo U1-T, a T is present immediately 5’ end of the cleavage site and base pairs with A (red type). The overlap displaces the 3’ T of U1-T, creating an unpaired 3’ flap. To ensure that 3’ flaps are held at 1 nt in length when oligo U1-T is used, a V (A, C, or G) is placed at the base of the 5’ flap, which does not pair with the indicated in A oligo 1-T (cyan arrow). Coloration of each sequence matches that used in Fig. 1a. Oligo sequences are listed in Supplementary Table 1, Sheet 1.

**Supplementary Fig. 2.**
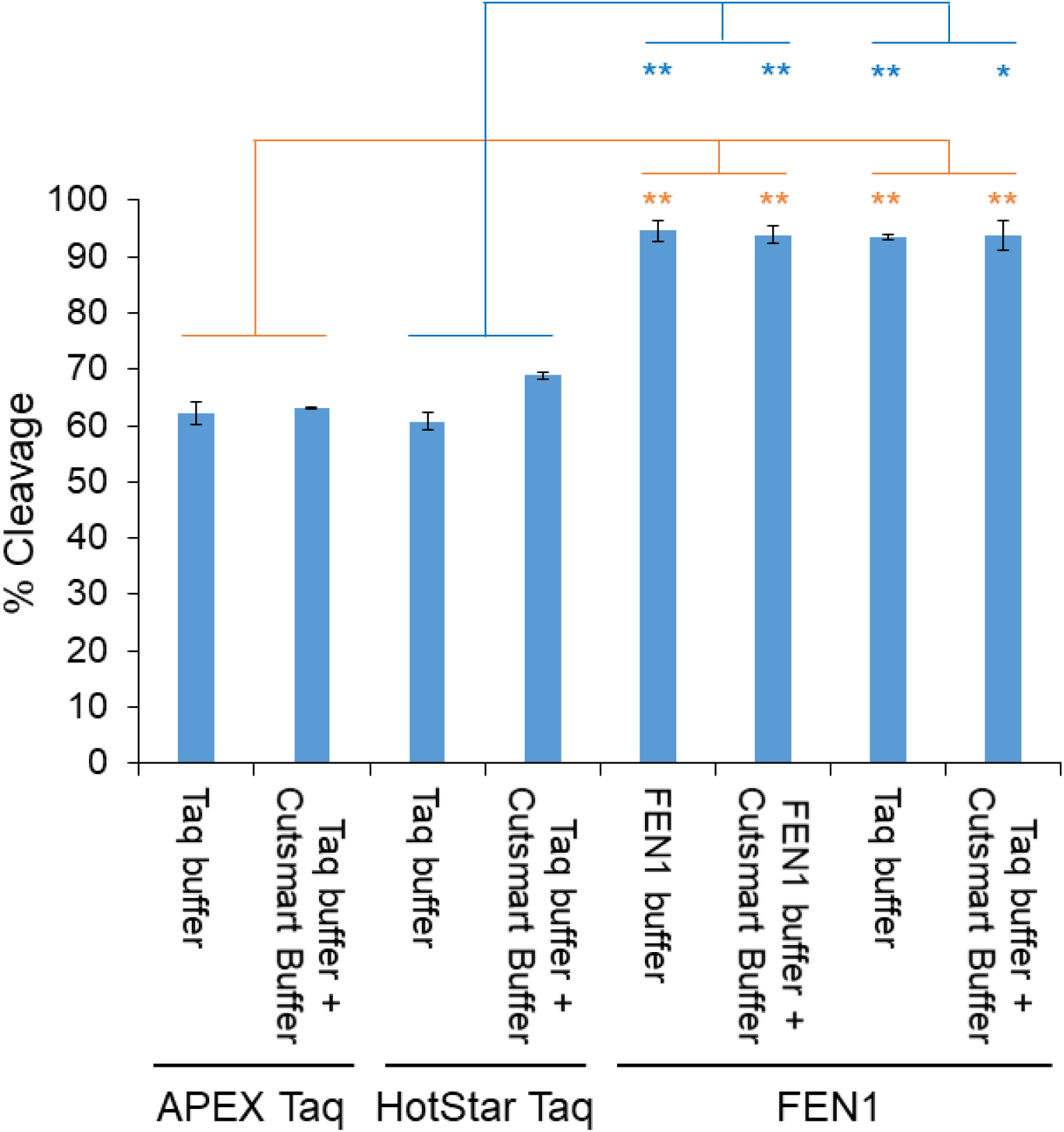
FEN cleavage efficiencies of Taq and FEN1. Percent cleavage of a 200-mer T proxy target assembled into a 129-nt 5’ flap by the indicated FENs using the indicated buffers. FEN1 cut the 200-mer with higher efficiency than Taq under all tested conditions (*n* = 2, mean ± range; paired Student’s *t* test, two sided, \**P* < .05, \*\**P* < .01). Oligos are described in Supplementary Table 1, Sheet 1.

**Supplementary Fig. 3.**
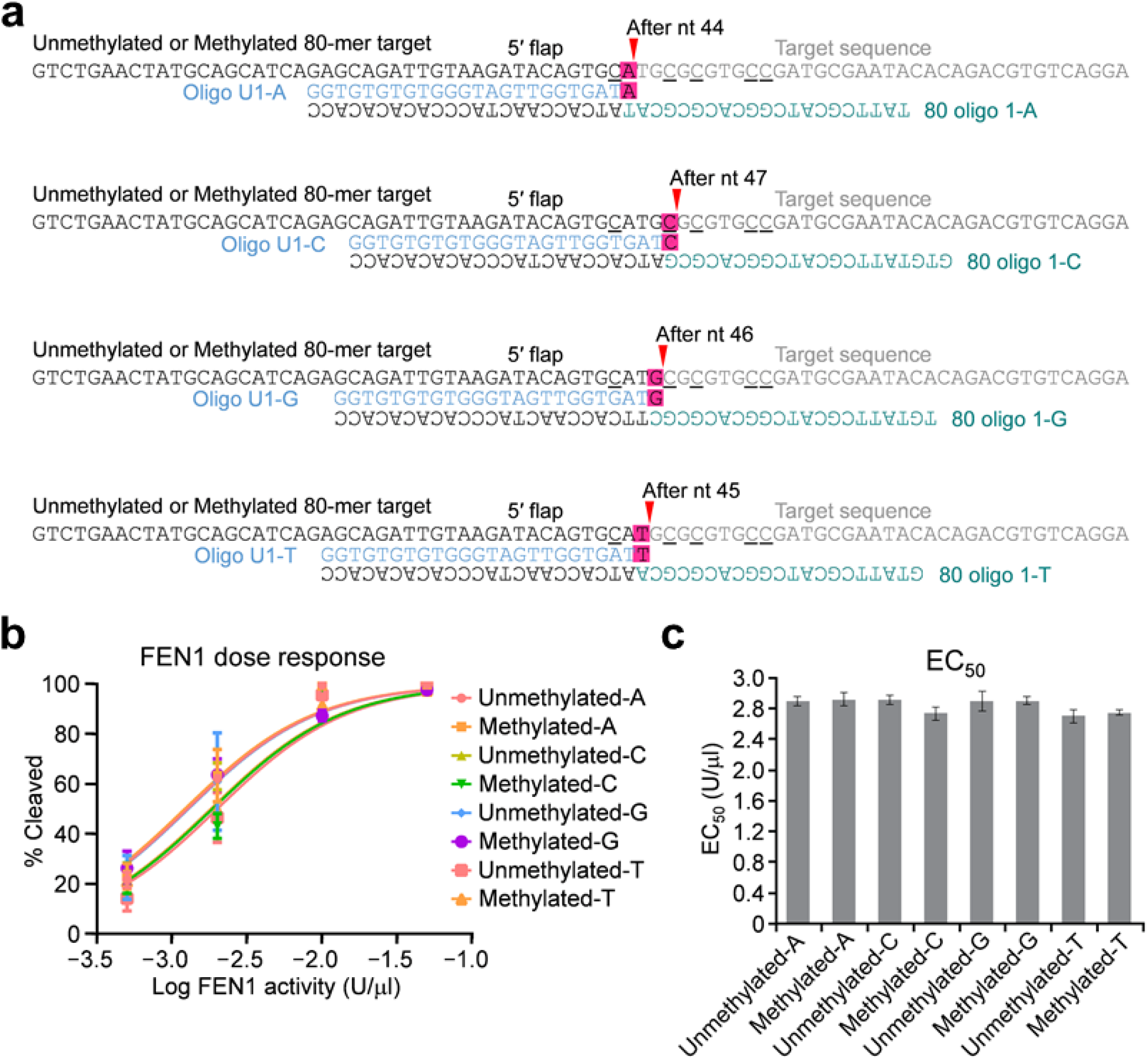
Equally efficient cleavage of double flaps with A, C, G, or T as ‘overlap’ in the absence and presence of nearby 5mC. **a**, Unmethylated and methylated double flap substrates used to test FEN1 5’ flap cleavage efficiency. Two 80-mers with the same sequence were synthesized without or with 5mC at positions 43, 47, 49, 53, and 54 (underlined). Both 80-mers were reconstituted into four different double flaps by incubation with their respective flap adapter comprised of oligo U1-N and its corresponding 80 oligo 1-N. The 1-nt ‘overlap’ that displaces the 3’-terminal nucleotide of each oligo U1-N to create an unpaired 1-nt 3’ flap is highlighted pink. Red arrowheads, FEN1 cleavage sites. All oligos are described in Supplementary Table 1, Sheet 1. **b,c**, Percentages cleaved of each 80-mer reconstituted into a -A, - C, -G, or -T double flap shown in **a** were quantified by qPCR and plotted as response *vs*. dose (**b**) and EC_50_ (**c**). A one-way ANOVA test of EC_50_ data in **c** showed no significant differences between any pair of conditions (*n* = 3, mean ± SD). Oligos are described in Supplementary Table 1, Sheet 1.

**Supplementary Fig. 4.**
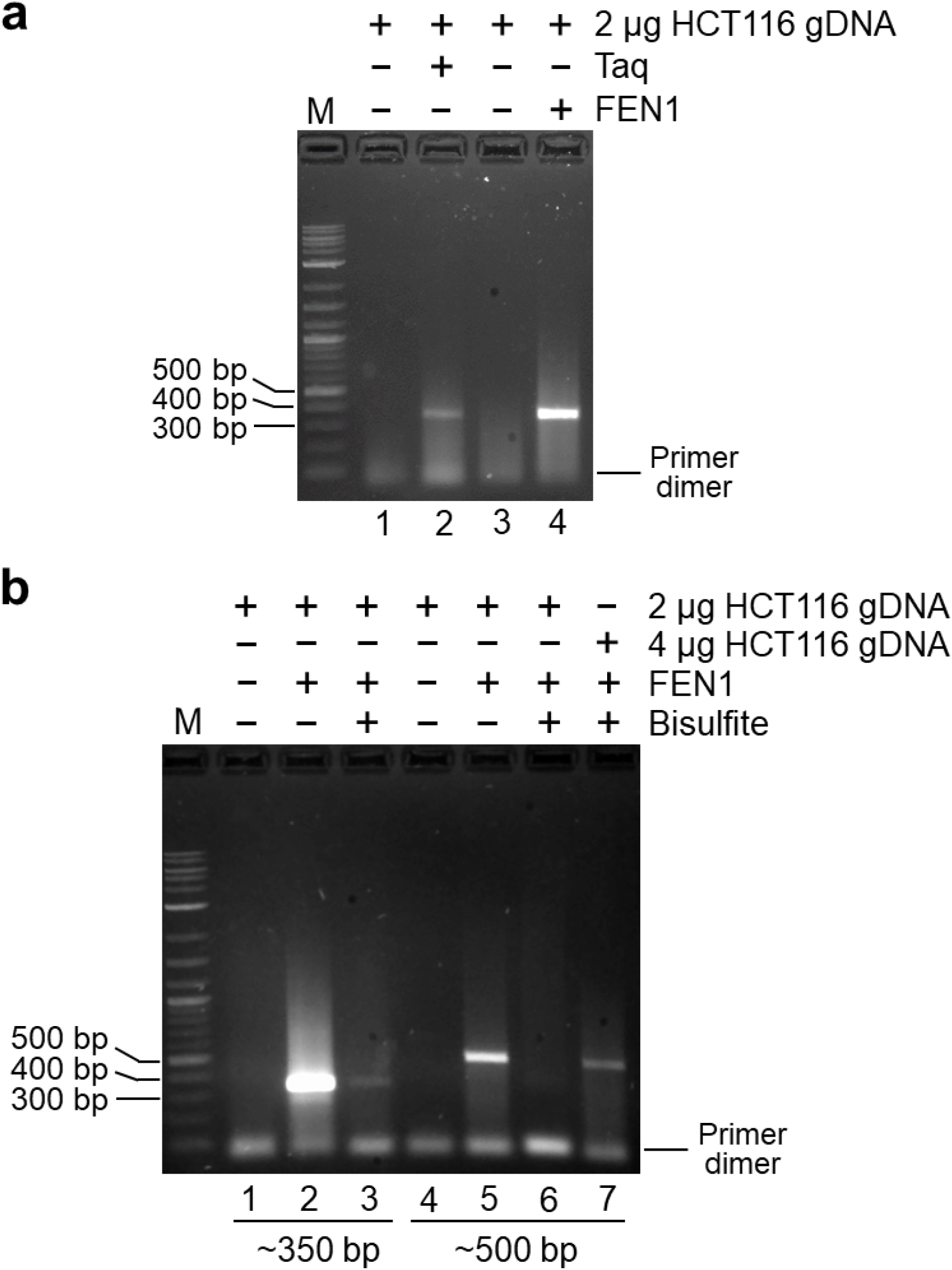
FEN activity-dependent enrichment of single-copy human sequences with and without bisulfite conversion. **a,b**, Human HCT116 gDNA (2 or 4 μg) was processed by FENGC with a 1-nt 3’ flap as described in Figure 1a, using flap adapters 1-T and 2-T specific for 11-promoter panels that excise ∼ 300-nt (**a**) or ∼ 300-nt and ∼ 450-nt (**b**) sequences in separate reactions. Both sets of reactions contained the same flap adapters 1-T, whereas the flap adapters 2-T in the ∼ 450-nt reactions in **b** were shifted downstream (Supplementary Table 1, Sheets 2 and 3; Supplementary Table 2, Sheets 1 and 2). Reactions also contained either no enzyme, 2 U Taq in 1x Taq buffer, or 32 U FEN1 in 1x FEN1 buffer as indicated. Enriched targets (post-exonuclease treatment and purification) were amplified by standard PCR in **a** or standard PCR (lanes 1, 2, 4, and 5) or BS-PCR (lanes 3, 6, and 7) in **b**. The crude amplification products, expected to be ∼350 bp and ∼500 bp, were electrophoresed on a 1% agarose gel. The results shown in both gels are representative of two independent experiments using the same HCT116 gDNA.

**Supplementary Fig. 5.**
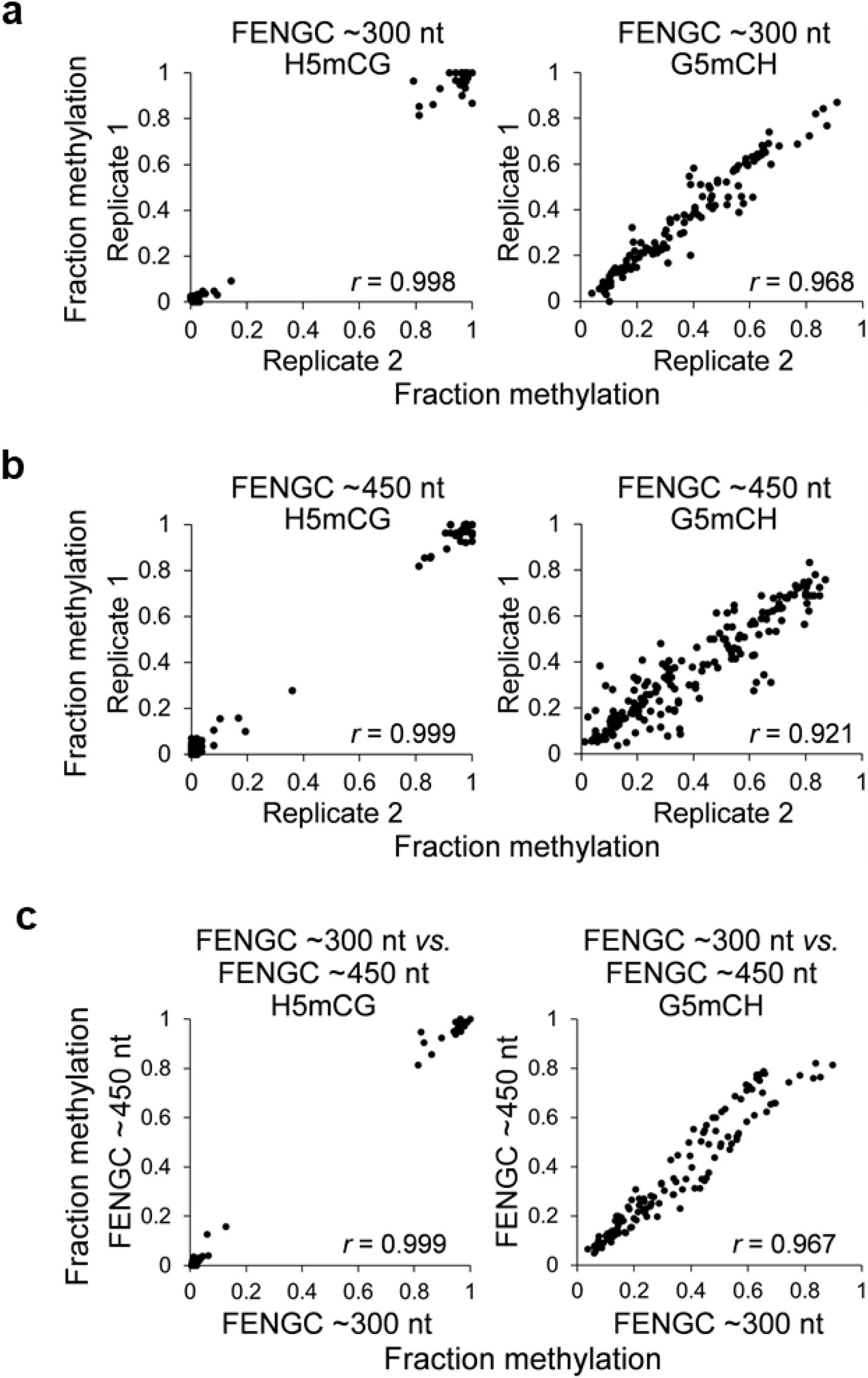
High reproducibility of MAPit-FENGC with BS-seq of different-size targets. **a,b**, Pearson correlation plots of H5mCG and G5mCH levels between two biological replicates of MAPit-FENGC, processing two independent cultures of M.CviPI-treated human GBM L0 using FEN1 to enrich for ∼ 300-nt (**a**) and ∼ 450 nt targets (**b**) from the 11-promoter panel (Supplementary Table 1, Sheets 2 and 3, respectively). **c**, Correlation plots of H5mCG and G5mCH levels between sequences in common between the ∼ 300-nt and ∼ 450 nt targets. Hi-Fi BS-seq CCS PacBio Sequel reads (≥ 5 sequencing passes with ≥95% conversion) from the two libraries for each condition were aligned to the reference sequences. Methylation levels are from 6 promoters with >40 aligned reads in each biological replicate (≥ 150x in combined replicates) in both the ∼300-nt and ∼450-nt targets (Supplementary Table 3).

**Supplementary Fig. 6.**
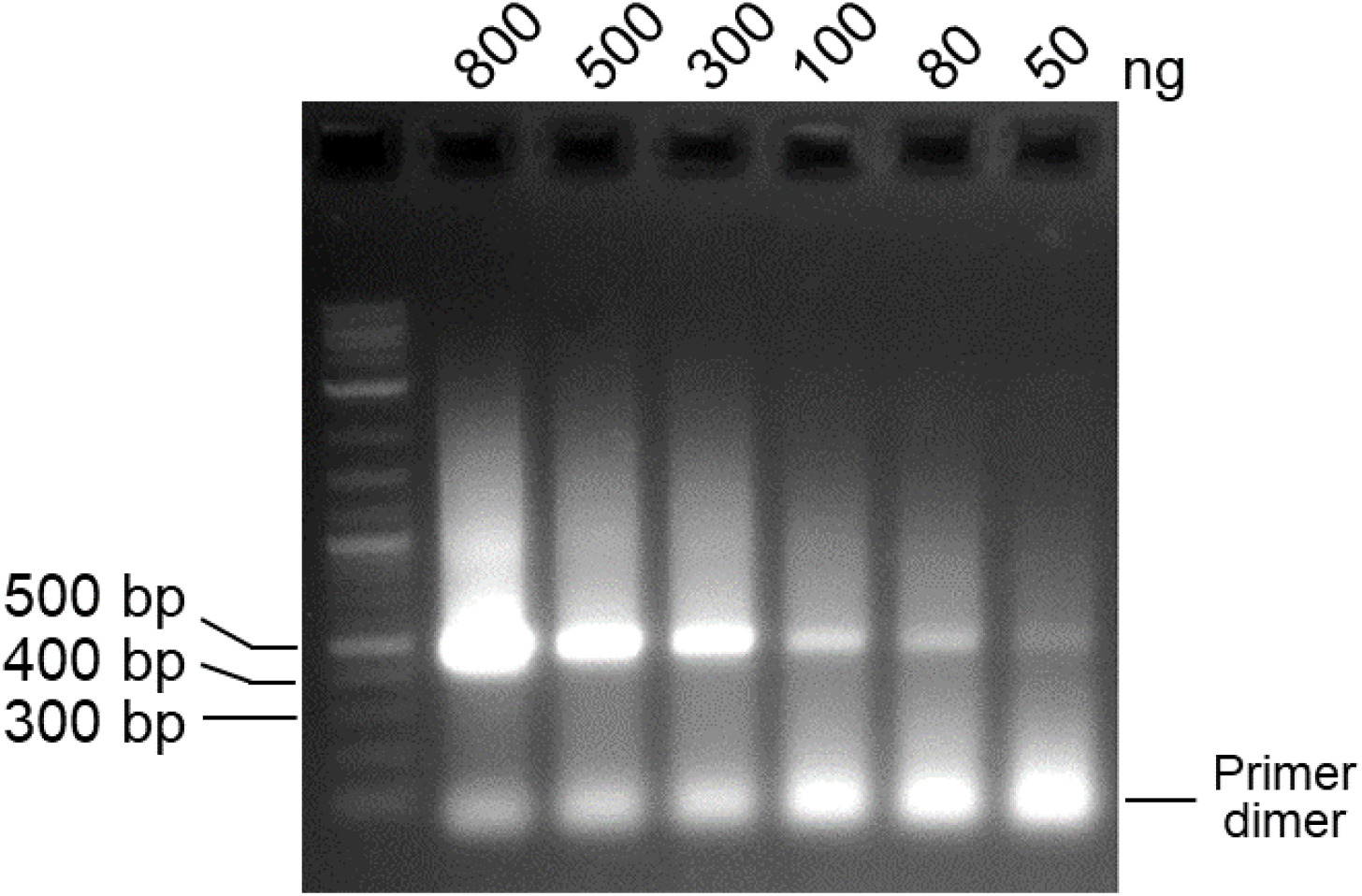
MAPit-FENGC with EM-PCR. Agarose gel electrophoresis of FENGC amplicon yields from the indicated amounts of M.CviPI-treated GBM L0 gDNA following fragmentation with mild sonication. The 5’ flaps of 119 promoter sequences of ∼ 450 nt (Supplementary Table 1, Sheet 3) were cleaved by 32 U FEN1 within double flap structures with 3’ flaps comprised of one T. Crude EM-PCR amplification products were electrophoresed without purification. For library construction, the primer dimer is readily removed by a single round of size selection with AMPure beads (data not shown). The shown results are representative of two independent experiments.

**Supplementary Fig. 7.**
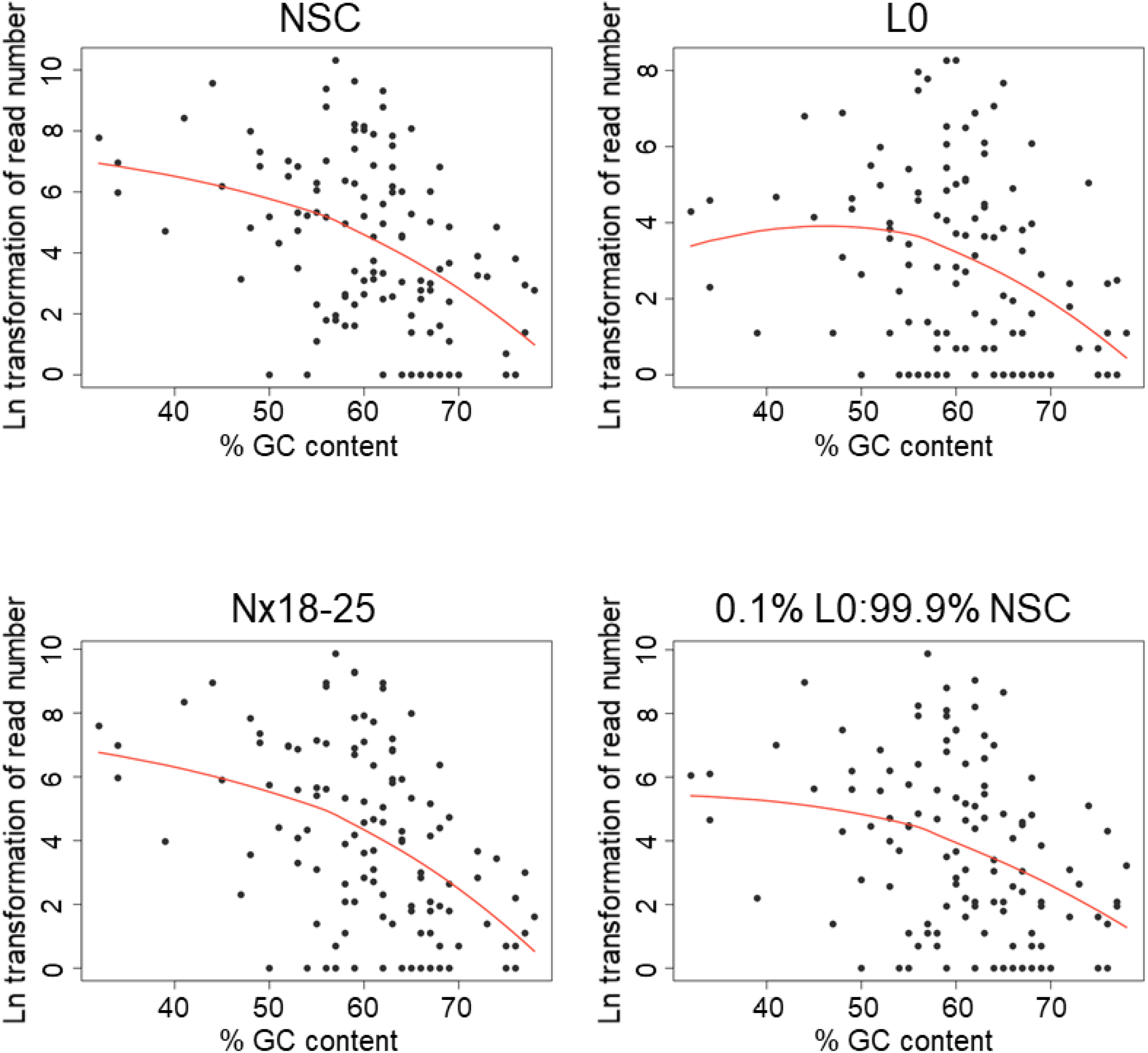
GC content exhibits a moderate, negative correlation with MAPit-FENGC CCS read number. LOESS fit curves of the natural logarithm transformation of the filtered read number in the combined biological replicates plus 1 *versus* GC content for each of the 119 targets of ∼ 450 nt for the indicated samples (Supplementary Table 5). Hi-Fi CCS reads from the 119-target MAPit-FENGC libraries were filtered for ≥95% HCH to HTH conversion and ≥ 95% coverage of the length of each reference sequence.

**Supplementary Fig. 8.**
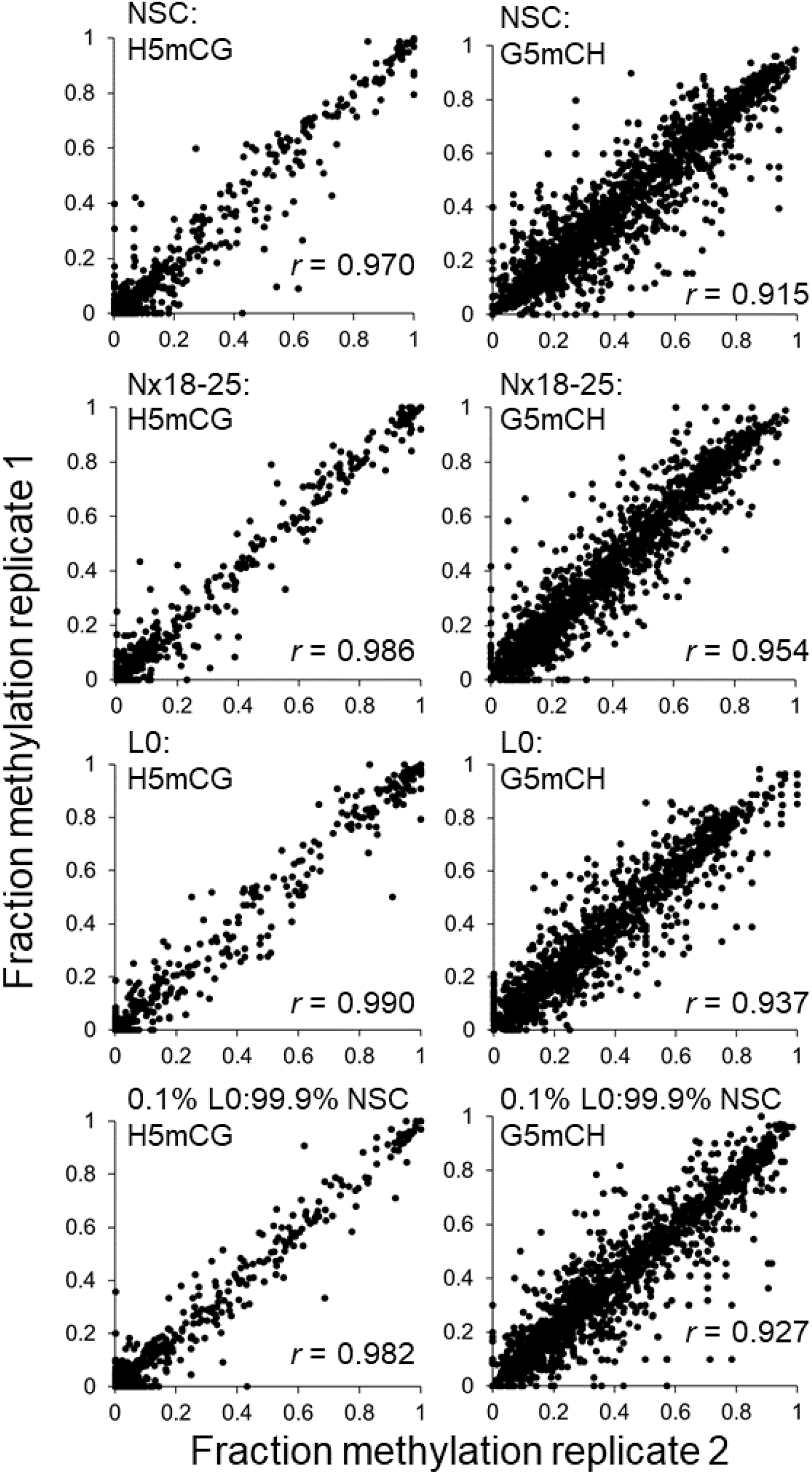
High reproducibility of MAPit-FENGC utilizing EM-PCR of a 119-promoter panel. Correlation plots between the fraction of methylation of each HCG and GCH site in the independent biological replicates of the indicated MAPit-FENGC samples enriched for the 119 human promoter sequences of ∼ 450 nt. Only targets with ≥ 10 Hi-Fi CCS reads aligned and filtered for ≥ 95% conversion and ≥ 95% coverage of reference sequence length in both replicates were included (Supplementary Table 5). This equated to 64 targets for NSC, 52 targets for L0, 64 targets for Nx18-25, and 63 targets for the 0.1% L0 gDNA:99.9% NSC gDNA mixtures.

**Supplementary Fig. 9.**
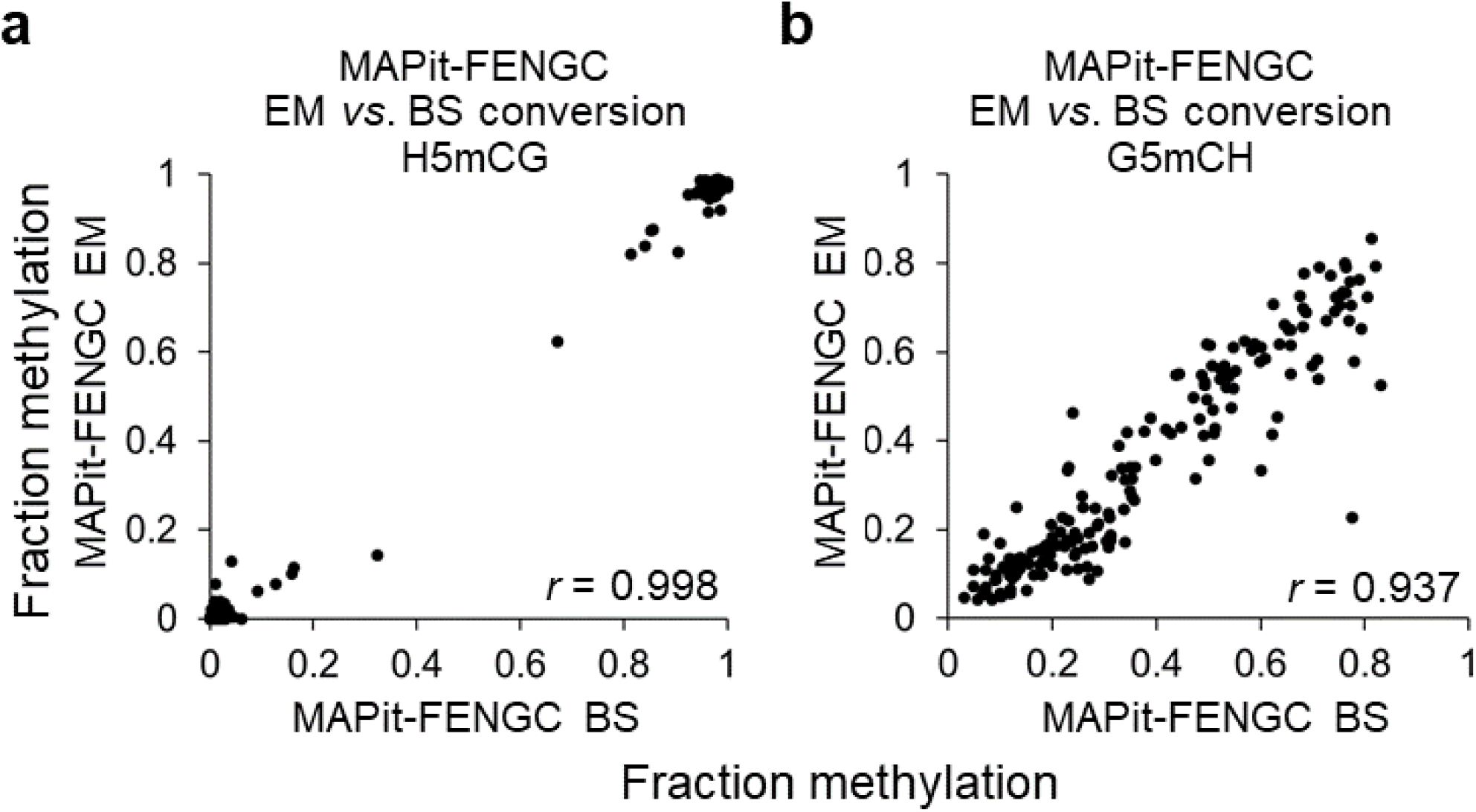
Strong correlation between MAPit-FENGC using EM and bisulfite conversion. **a,b**, Correlation plots of the fractions of methylation at each HCG (**a**) and GCH site (**b**). Data plotted for the 6 targets of ∼ 450 nt with ≥ 36 aligned and filtered CCS reads in the combined GBM L0 duplicates for EM conversion (Supplementary Table 5) and bisulfite conversion (Supplementary Table 3).

**Supplementary Fig. 10.**
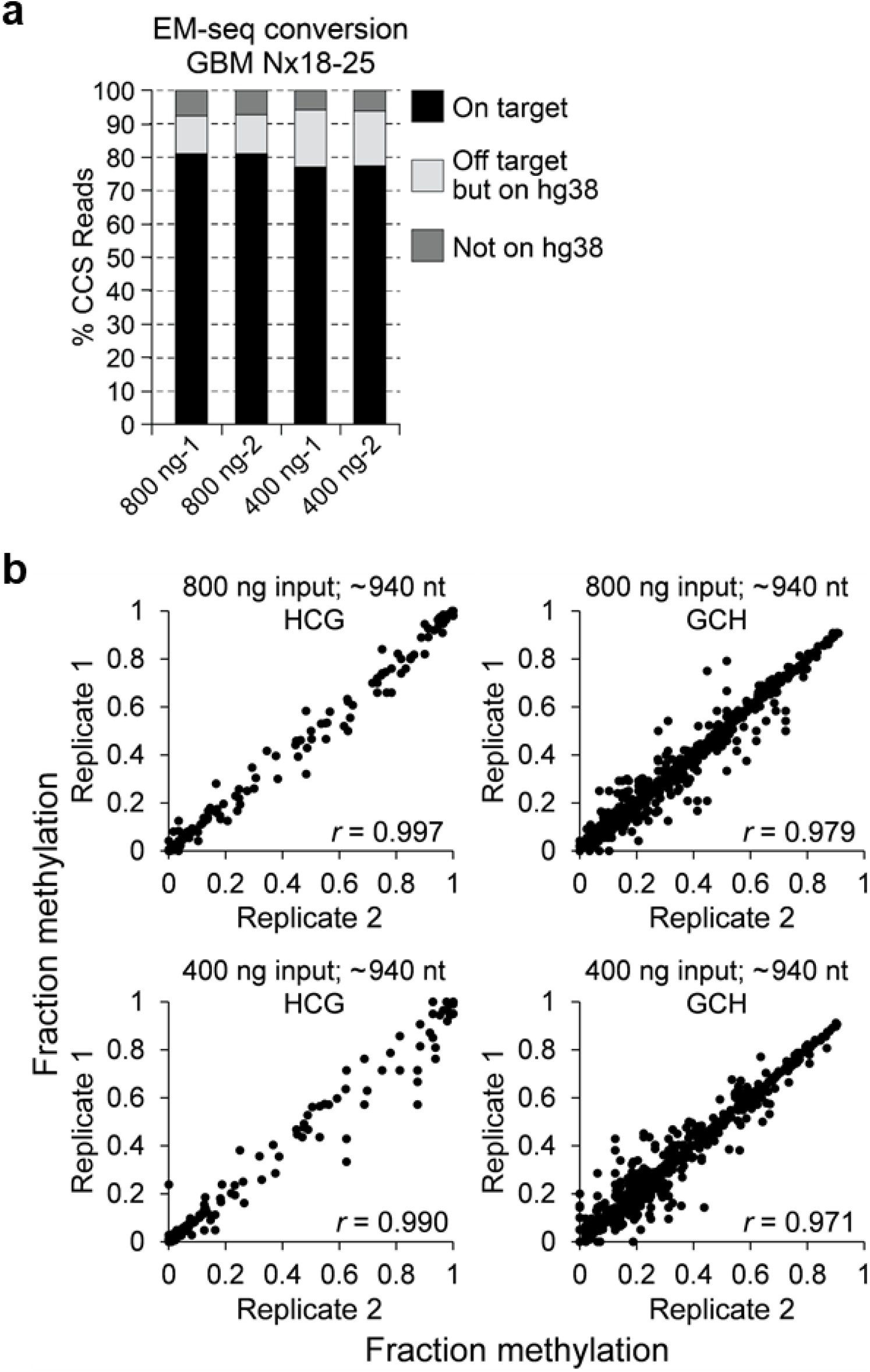
Performance of MAPit-FENGC for captured ∼ 940-nt targets with two different gDNA amounts. **a**, Proportions of Hi-Fi CCS reads for MAPit-FENGC of 45 promoter sequences (Supplementary Table 1, Sheet 4; Supplementary Table 2, Sheet 3) mapping to or off the human genome for the indicated sequenced libraries (Supplementary Table 8). **b**, Correlation plots of the fraction of methylation of each HCG and GCH motif in the independent biological replicates of the indicated MAPit-FENGC samples. Only targets with ≥ 14 Hi-Fi CCS reads aligned and filtered for ≥ 95% conversion and ≥ 95% coverage of reference sequence length in both replicates were included (Supplementary Table 9). This equated to 13 and 12 targets for inputs of 800 ng and 400 ng, respectively.

**Supplementary Fig. 11.**
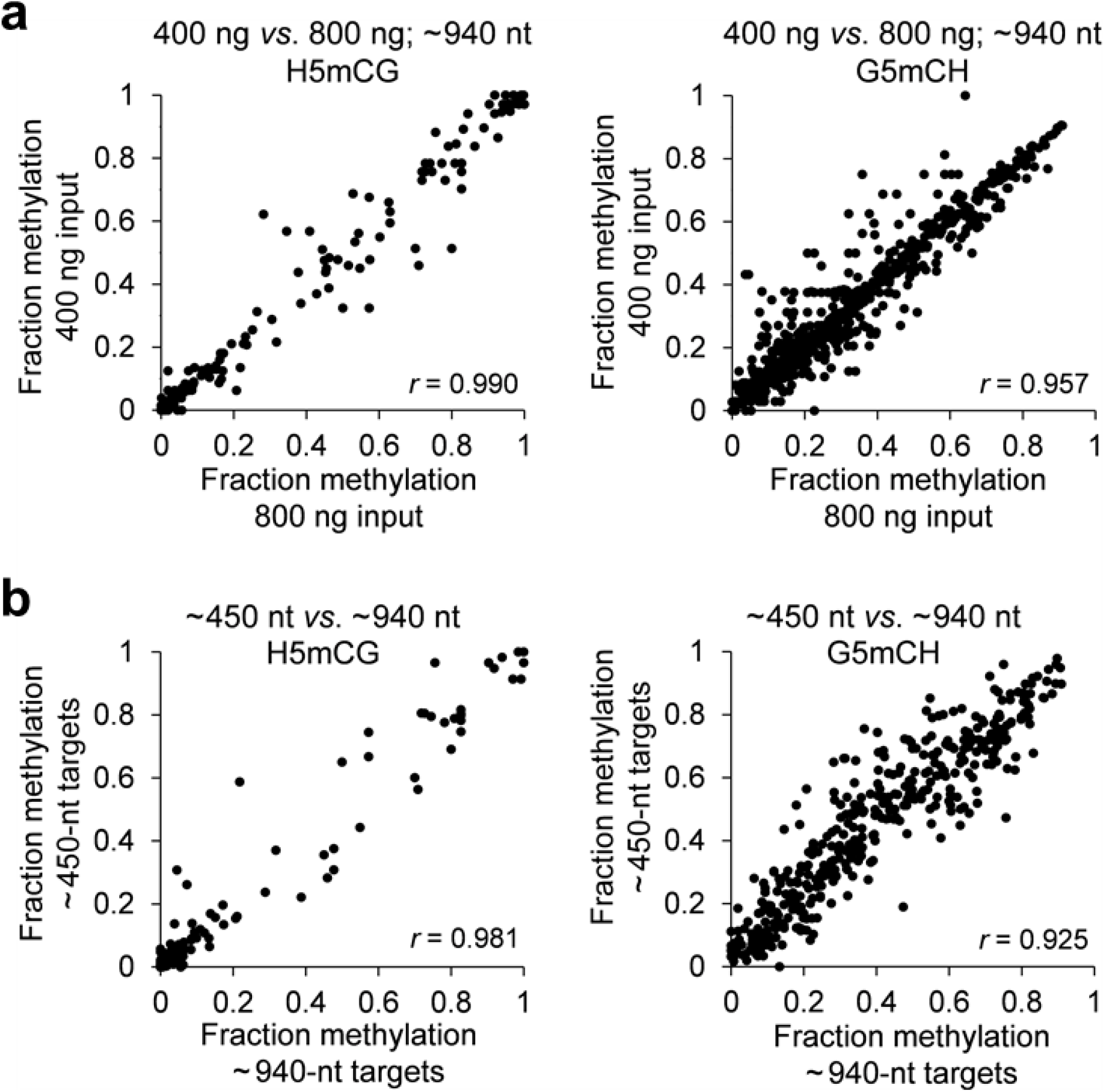
Performance of MAPit-FENGC for captured ∼ 940-nt targets with two different gDNA amounts and comparison to ∼ 450-nt targets. **a,b**, Correlation plots of the fraction HCG and GCH methylation in the combined biological replicates of ∼ 940-nt targets for 400 *vs*. 800 ng input gDNA (**a**) and for sequences in common between ∼ 940-nt *vs*. ∼ 450-nt targets (**b**) from M.CviPI-treated Nx18-25. After filtering HiFi CCS reads for ≥ 95% conversion and ≥ 95% coverage of reference sequence length (Supplementary Table 9), methylation levels of 13 targets with ≥ 15x coverage in the combined biological replicates were plotted.

**Supplementary Fig. 12.**
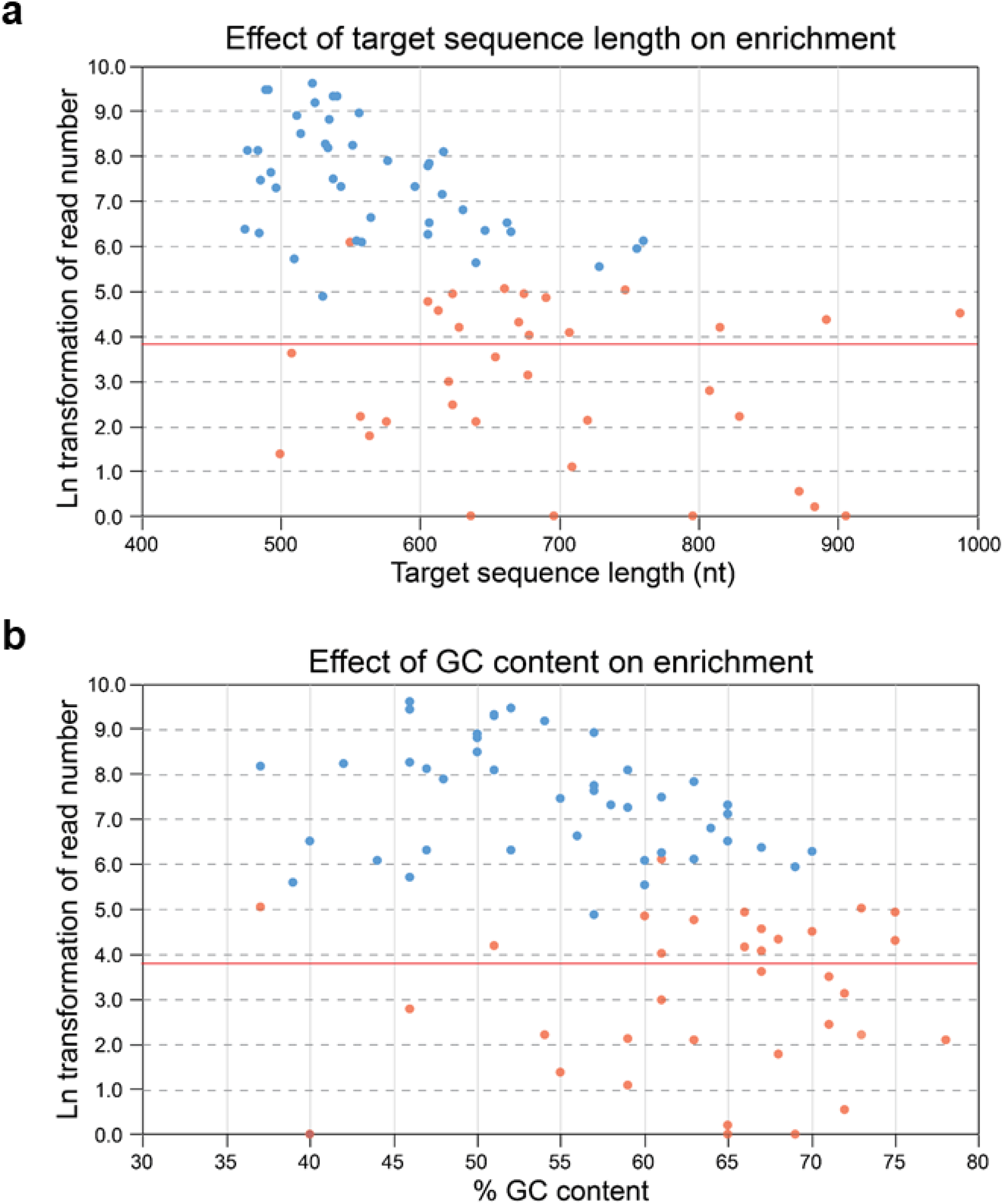
Effect of target sequence length and GC content on performance of MAPit-FENGC with EM-seq on primary mouse monocytes. **a,b**, Natural logarithm transformation of filtered CCS read number + 1 *versus* (**a**) target length and (**b**) % GC content for the 78-target murine gene panel enriched from gDNA isolated from bone marrow-derived monocytes probed with M.CviPI (*n* = 4 female mice). In (**a**) and (**b**), blue and red data points represent the combined bioreplicates of 43 targets included and 35 excluded, respectively, from the statistical analysis of Supplementary Table 12. Target loci were excluded because their read number fell below the 100-read per sample threshold (red horizontal line) or presence of too many duplicates, typically observed for targets > 760 nt or > 70% GC content.

## SUPPLEMENTARY TABLES

**Supplementary Table 1**. FENGC oligonucleotides used in this study.

**Supplementary Table 2**. Characteristics of FENGC targets used in this study.

**Supplementary Table 3**. CCS reads aligned to 11 human targets of ∼ 300 nt *vs*. ∼ 450 nt used in FENGC assay development with standard PCR *vs*. BS-PCR.

**Supplementary Table 4**. CCS reads on- and off-target for 119 ∼ 450-nt human targets in MAPit-FENGC.

**Supplementary Table 5**. Filtered CCS reads aligned to 119 ∼ 450-nt human targets in MAPit-FENGC using EM-seq.

**Supplementary Table 6**. Differential H5mCG and G5mCH in NSC and GBM Nx18-25 among ∼ 450-nt targets determined by MAPit-FENGC using EM-seq.

**Supplementary Table 7**. SNPs and indels detected by FENGC using standard PCR of 119 ∼ 450-nt human targets.

**Supplementary Table 8**. CCS reads on- and off-target for 45 ∼ 940-nt human targets in MAPit-FENGC using EM-seq.

**Supplementary Table 9**. Filtered CCS reads aligned to 45 ∼ 940-nt human targets in MAPit-FENGC using EM-seq.

**Supplementary Table 10**. CCS reads on- and off-target for 78 ∼ 620-nt mouse targets in MAPit-FENGC using EM-seq.

**Supplementary Table 11**. Filtered CCS reads aligned to 78 ∼ 620-nt mouse targets in MAPit-FENGC using EM-seq.

**Supplementary Table 12**. Differential H5mCG and G5mCH determined by MAPit-FENGC using EM-seq of ∼ 620-nt mouse targets.

**Supplementary Table 13**. FENGC oligonucleotide concentrations for different numbers of targets.

## Notes

### Competing Interest Statement

Mingqi Zhou, Nancy Nabilsi, and Michael Kladde are inventors (assignee, University of Florida) of pending patent application (PCT/US2022/020624, filed March 16, 2022) for the FENGC technology.

### Summary of Updates

Minor textual updates, e.g., added additional Methods; uploaded Supplementary Tables to Excel files (to replace Pdf versions); updated Distribution/Reuse terms

